# The distinct role of ALDH1A1 and ALDH1A3 in the regulation of prostate cancer metastases

**DOI:** 10.1101/2021.05.08.443223

**Authors:** Ielizaveta Gorodetska, Anne Offermann, Jakob Püschel, Vasyl Lukiyanchuk, Diana Gaete, Anastasia Kurzyukova, Vera Labitzky, Franziska Schwarz, Tobias Lange, Franziska Knopf, Ben Wielockx, Mechthild Krause, Sven Perner, Anna Dubrovska

**Affiliations:** OncoRay-National Center for Radiation Research in Oncology, Faculty of Medicine and University Hospital Carl Gustav Carus, Technische Universität Dresden and Helmholtz-Zentrum Dresden-Rossendorf, Dresden, Germany; Institute of Pathology, University Hospital Schleswig-Holstein, Luebeck, Germany; Pathology, Research Center, Borstel, Leibniz Lung Center, Borstel, Germany; Institute of Clinical Chemistry and Laboratory Medicine, Technische Universität Dresden, Dresden, Germany; Technische Universität Dresden, CRTD - Center for Regenerative Therapies TU Dresden, Center for Healthy Aging, Dresden, Germany; Institute of Anatomy and Experimental Morphology, Center for Experimental Medicine, University Cancer Center, Hamburg, University Medical Center Hamburg-Eppendorf, Germany; Department of Radiotherapy and Radiation Oncology, Faculty of Medicine and University Hospital Carl Gustav Carus, Technische Universität Dresden, Dresden, Germany; National Center for Tumor Diseases (NCT), partner site Dresden: German Cancer Research Center (DKFZ), Heidelberg; Faculty of Medicine and University Hospital Carl Gustav Carus, Technische Universität Dresden, and Helmholtz-Zentrum Dresden-Rossendorf (HZDR), Dresden, Germany; Helmholtz-Zentrum Dresden-Rossendorf, Institute of Radiooncology-OncoRay, Dresden, Germany; German Cancer Consortium (DKTK), partner site Dresden and German Cancer Research Center (DKFZ), Heidelberg, Germany

**Keywords:** prostate cancer, bone metastases, cancer stem cells, aldehyde dehydrogenase

## Abstract

Cancer stem cells (CSC) are characterized by high self-renewal capacity, tumor-initiating potential, and therapy resistance. Aldehyde dehydrogenase (ALDH)+ cell population serves as an indicator of prostate CSCs with increased therapy resistance, enhanced DNA double-strand break repair, and activated epithelial-mesenchymal transition (EMT) and migration. Numerous ALDH genes contribute to ALDH enzymatic activity; however, only some of them showed clinical relevance. We found that ALDH1A1 and ALDH1A3 genes functionally regulate CSC properties and radiation sensitivity of PCa. We revealed a negative correlation between ALDH1A1 and ALDH1A3 expression in publicly available prostate cancer (PCa) datasets and demonstrated that ALDH1A1 and ALDH1A3 have opposing predictive value for biochemical recurrence-free survival. Our data suggest an association of ALDH1A1 with the metastatic burden, elucidating the role of ALDH genes in the metastatic spread and homing to the bone, which can be, at least partially, attributed to regulating the transforming growth factor beta 1 (TGFB1) and matrix metalloproteinases (MMPs). ALDH genes play a diverse role in PCa development under AR and β-catenin-dependent regulation, with ALDH1A1 becoming dominant in later stages of tumor development when PCa cells gain androgen independence. Taken together, our results indicate that ALDH1A1 and ALDH1A3 modulate PCa radiosensitivity, regulate CSCs phenotype, and spread of PCa cells to the bone, therefore having clinical implication for identifying patients at high risk for progression to metastatic disease.

## Introduction

Prostate cancer (PCa) is the second most commonly diagnosed malignancy in men, accounting for 1.3 million new cases worldwide in 2018 (Bray et al., 2018). At early stage, PCa may be permanently controlled with surgery or radiotherapy alone or together with androgen deprivation therapy. However, bone metastatic PCa has a heterogenic intra- and interpatient response to treatment (Morin et al., 2017). Although the 5-year survival rate for patients with localised disease is around 99%, metastatic PCa often remains incurable and is one of the primary causes of cancer-related death in men (Bray et al., 2018). PCa is a complex disease with heterogeneous clinical behaviour (Skvortsov et al., 2018). Although of a high clinical importance, the current prognostic factors such as TNM staging, the Gleason scoring and the prostate-specific antigen (PSA) levels do not explain substantial variability in the treatment outcomes that goes beyond the clinical-pathological parameters. While many prognostic signatures for PCa were recently developed, they are often confounded by clonal heterogeneity and multifocal tumor growth (Brastianos et al., 2020). Identification of prostate cancer stem cell (CSC) populations, which maintain primary and metastatic tumor growth offered new opportunities for the biomarker development and therapeutic intervention. CSCs are characterised by high self-renewal capacity, tumour-initiating potential and therapy resistance (Peitzsch et al., 2019). Nowadays, several markers allow to identify and isolate prostate CSCs from patient-derived tumors and cell lines (Skvortsov et al., 2018). Nonetheless, the clinical application of distinct CSC subpopulations as prognostic indicators in PCa is still uncertain.

Aldehyde dehydrogenase (ALDH) is a family of metabolic enzymes responsible for the oxidation of cellular aldehydes to the corresponding carboxylic acids (Vassalli, 2019). High levels of ALDH activity measured by conversion of bodipy-aminoacetaldehyde (BAAA) into bodipy-aminoacetate (BAA) have been reported in CSCs of the different tumor types, including PCa. Therefore, this functional assay is widely used for identification and isolation of cancer cells with stem cell properties (Hoogen et al., 2010; Magnen et al., 2013; Nishida et al., 2012; Zhou et al., 2019). ALDH^+^ cell population serves as an indicator of prostate CSCs with increased therapy resistance, enhanced DNA double-strand break repair as well as activated epithelial-mesenchymal transition (EMT) and migration (Cojoc et al., 2015; Peitzsch et al., 2016). To date, scientists identified 19 ALDH isoform genes in the human genome, but not all contribute to ALDH enzyme activity (Vasiliou and Nebert, 2005; Zhou et al., 2019). Accordingly, it is reasonable to suggest that ALDH genes contributing to ALDH activity might play a role in regulating CSC populations. Identification of ALDH isozymes responsible for ALDH activity and CSC phenotype, and investigation of the molecular pathways by which CSCs cells survive radiotherapy and metastasize might contribute to the PCa biomarker development.

*ALDH1A1* has long been regarded as the main gene responsible for the ALDH^+^ activity measured by Aldefluor assay (Zhou et al., 2019). Nevertheless, other ALDH isoforms contributing to the regulating ALDH activity in CSCs could also play a role in tumor suppression or progression in a cancer-type dependent manner, either positively or negatively influencing the patient’s treatment outcome (Chang et al., 2018; Zhang et al., 2011; Zhou et al., 2019).

Our study aims to investigate the cellular processes and molecular mechanisms regulated by ALDH proteins, which contribute to PCa stemness and radioresistance. We hypothesise that *ALDH1A1* and *ALDH1A3* maintain the CSC phenotype and regulate bone metastasis-initiating cells and their survival after radiotherapy. We also validated ALDH1A1 and ALDH1A3 as potential biomarkers of clinical outcome and metastases on the cohort of PCa patients.

## Materials and Methods

### Patient’s samples

For the evaluation of ALDH1A1 and ALDH1A3 expression in human PCa, 33 benign prostatic samples, 457 primary PCa samples obtained by radical prostatectomy, 55 local recurrent or locally advanced PCa samples obtained by transurethral resection of the prostate, 35 lymph node metastases and 57 distant metastases were used for analysis. Benign samples, primary PCa samples and local recurrent or locally advanced tumors are from patients diagnosed with PCa in the Hospital of Goeppingen, Germany between 1997 and 2014. Lymph node and distant metastases are from patients treated in the University Hospital Schleswig-Holstein, Campus Luebeck between 2002 and 2015. Disease recurrence was defined as rising serum PSA level after radical prostatectomy indicating disease progression. Clinical material was collected with informed consent from all subjects. Ethical approval for this retrospective analysis of clinical and biological data was obtained from the local Ethics Committee.

### Immunohistochemistry

ALDH1A1 and ALDH1A3 protein expression was detected and quantified using immunohistochemistry (IHC). Therefore, fresh frozen paraffin embedded (FFPE) tissue blocks (donor blocks) were used to create tissue microarrays (TMA). Three representative cores per sample from donor blocks were placed into a TMA recipient using a semiautomated tissue arrayer (Beecher Instruments, Sun Prairie, WI, USA). Immunohistochemistry was performed after deparaffinization, following treatment with a primary anti-ALDH1A1-antibody (Santa Cruz Biotechnology, sc-374076) or anti-ALDH1A3-antibody (Atlas Antibodies, HPA046271) on the Ventana BenchMark (Roche, Basel, Switzerland) by using the IView DAB Detection Kit. Expression levels were evaluated by two pathologists (AO, SP) and categorised according to negative, low to moderate and high staining intensity. The expression of all three replicates per sample was considered, and highest expression in a single core was used for further analysis in cases of heterogeneous staining levels. Androgen receptor (AR) expression was detected and evaluated as described before (Becker et al., 2020).

### Cell lines and culture condition

PCa cell lines PC3 (derived from bone metastasis), LNCaP (derived from lymph node metastasis) and LNCaP-C42B (further named C42B, bone metastatic derivative subline of human prostate cancer LNCaP cell line) were purchased from the American Type Culture Collection (Manassas, VA, USA) and cultured according to the manufacturer’s recommendations in a 37°C incubator in an atmosphere with 5% CO_2_. PC3 cell line was cultivated in DMEM (Sigma-Aldrich); LNCaP and C42B cells in RPMI1640 medium (Sigma-Aldrich) supplemented with 10% FBS (PAA Laboratories) and 1 mM L-glutamine (Sigma-Aldrich). Radioresistant (RR) cell lines were established as described before (Cojoc et al., 2015; Peitzsch et al., 2016). Cells radioresistance was verified by radiobiological clonogenic cell survival assay. Corresponding age-matched non-irradiated parental cells were used as controls for RR cell lines.

The murine prostate carcinoma cell line RM1 bone metastatic (BM) was established by continual intracardiac injection of RM1 cells transfected with the GFP-expressing plasmid (RM1-GFP) into C57BL/6 mice followed by isolation of cells from bone tumors as was described previously (Power et al., 2009). RM1(BM) cells were cultured in RPMI1640 medium (Sigma-Aldrich) supplemented with 10% FBS (PAA Laboratories) and 1 mM L-glutamine (Sigma-Aldrich).

### Establishment of color-coded PC3 cell lines

Color-coding of PC3 cell line with green or red fluorescent proteins was done with pWPXL vector and its derivative construct where EGFP was replaced by tdTomato. HEK293 cells were transfected by these constructs along with psPAX2 and pMD2.G plasmids using calcium phosphate method to produce replication incompetent lentiviral particles. Supernatant from transfected cells was collected for 3 days, pooled, cleared through 0.45um filter and applied on PC3 cells overnight. Transduced PC3 were passaged twice to expand and eliminate any residual lentivirus and after that, populations stably expressing corresponding fluorescent proteins were isolated via fluorescence-activated cell sorting (FACS). To obtain purified cell populations, GFP^+^ or tdTomato^+^ cells were isolated using FACS on the BD FACSAria™ III (BD Biosciences, Becton, Dickinson and Company). For this, the cells were detached using Accutase (PAA), taken up in Flow buffer (DPBS (Sigma Life Science) supplemented with 5% FBS, 1% HEPES (Sigma Life Science), 1mM EDTA (Sigma-Aldrich)), and stained with 7-aminoactinomyocin D (7AAD; Sigma Life Science) to exclude dead cells. The final purity of the sorted cell populations was 88.2% for GFP^+^ cells and 83.3% for tdTomato^+^ cells based on the reanalysis.

### shRNA-mediated gene silencing

RM1(BM) cells were transfected with pLKO.1 puro vector constructs expressing shRNA against mouse *Aldh1a1* or *Alhd1a3* and non-silencing control shRNA using Lipofectamine 2000 Transfection Reagent (Thermo Scientific, Waltham, MA) according to the manufacturer’s instructions. First, 3×32^5^ cells were seeded into individual wells of 6-well plates. For each well, 3 μg shRNA plasmid DNA and 13μl of Lipofectamine 2000 were diluted in 250 μl volumes of Opti-MEM reduced serum medium (Invitrogen) and mixed gently. After 5 minutes of incubation at room temperature, the DNA and the Lipofectamine 2000 complex was formed and then added to each well containing cells and medium. 48 hours after transfection, the Aldh1a1- or Aldh1a3-expressing cells were selected with puromycin at a concentration of 4 μg/ml. The list of shRNA constructs is represented in Supplementary table 1.

### siRNA-mediated gene silencing

The cells were grown until confluency of 60-80% in complete medium. According to the size of the well and manufacturer’s instructions, the Lipofectamine RNAiMAX (Thermo Scientific, Waltham, MA) and siRNAs were diluted at the corresponding concentrations in Opti-MEM reduced serum medium. The diluted Lipofectamine and siRNA were then gently mixed 1:1 in a tube and incubated for 5 minutes at room temperature. The corresponding volume of the mixture was added to each well, rocking the plates back and forth and the cells were incubated at 37°C in a CO^2^ incubator for 48 h. Cells transfected with unspecific siRNA (scrambled siRNA or siSCR) were used as a negative control in all knockdown experiments. The siRNA target sequences were obtained from the Life Technologies website and corresponding RNA duplexes were synthesized by Eurofins. The siRNA oligos used in the study are represented in Supplementary table 2.

### PCa cell preparation for injection into 2 days post fertilization (dpf) Zebrafish

PC3 color-coded cells were trypsinazed (0.25 % trypsin-ethylenediaminetetraacetic acid, Gibco) with subsequent addition of growth media to stop the reaction. The cell suspension was then centrifuged at 1200 rpm for 5 minutes and pellet was resuspended in PBS to achieve 1×32^6^ cells/ml. The suspension was transferred into a 1.5 ml Eppendorf tube and centrifuged at 800 rpm for 10 minutes. Afterwards, cancer cells were washed one time in PBS and one time in Tx Buffer (PBS, 1% Penicillin/Streptomycin, 1.5 μM EDTA). The pellet of 1×32^6^ cells was resuspended in 10 μl Tx Buffer.

### Injection of PCa cells into 2dpf Zebrafish

Adult zebrafish of strain *flk1:CFP (Tg(kdrl:CFP)zf410Tg)* (Hess and Boehm, 2012) were incrossed for the generation of embryos with a vessel marker. Subsequently, eggs were collected, selected for CFP^+^ signal using stereomicroscope (Leica MZ16 FA), transferred to 10cm plastic dish and kept in E3-medium (5 mM NaCl, 0.17 mM KCl, 0.33 mM CaCl2, 0.33 mM MgSO4) at 28°C in incubator (TS608/2-1, WTW). 2dpf embryos were mechanically dechorionated with sharp tweezers and anesthetized with 0.02% Tricaine solution (MS222, Merck) in plastic dish. The larvae were transferred on a grooved 1.5% agarose bed casted prior to experiments. Fine borosilicate glass tubes with filaments were pulled into injection capillaries by Flaming/Brown Micropipette Puller (Model P-97, Sutter Instruments Co.). Capillary diameter was adjusted to 20μm by cropping it with a tweezer. Cells were mixed immediately prior to injection again by repeated pipetting. 6μl of cell suspension were loaded into capillary, which in turn was inserted into microinjector (MM3301R, Marzhauser). Final injection volume was adjusted to 4nl (to introduce around 400 cells in total) and cells were injected into the Duct of Cuvier (DoC) of anesthetized zebrafish larvae with the help of a pneumatic pump (Pneumatic PicoPump PV 820, WPI) and binocular (SZX10, Olympus). The fish were transferred to new 10cm dish containing fresh E3-medium at a density of up to 50 larvae per dish and incubated at 33°C until 5dpf.

### Imaging of injected Zebrafish larvae

5dpf, 1 ml of 1% low melting point agarose (Biozym Scientific) aliquots were melted in water bath at 68°C. Upon melting, tubes were transferred to 42°C water bath and 20 μl of 0.4% Tricaine solution were added per aliquot. Meanwhile, zebrafish larvae were anesthetized with 0.02% Tricaine solution added to 10cm dish containing E3-medium. Immersed in agarose, injected larvae were transferred on microwell dishes with transparent glass window (35mm with 14 mm microwell, MatTek Corporation). After solidification of agarose, E3-medium with 0.01% Tricaine was carefully added until the agarose was fully submerged. Subsequently, tail region of fish was imaged by Dragonfly Spinning Disc Confocal Microscope (Andor Technology). Extravasation and survival were assessed in Fiji software.

### Clonogenic cell survival assay

Radiobiological clonogenic assay was performed as described previously (Cojoc et al., 2015; Peitzsch et al., 2016). Cells were plated at a density of 1000-4000 cells/well depending on the cell line and treatment in 6-well plates in triplicates. Next day cells were irradiated with doses of 2, 4 and 6 Gy of X-rays (Yxlon Y.TU 320; 200 kV X-rays, dose rate 1.3Gy/min at 20 mA) filtered with 0.5 mm Cu. Absorbed dose was measured using a Duplex dosimeter (PTW). Cells were incubated in a humidified 37°C incubator supplemented with 5% CO^2^ that allow them to form colonies. 10 days later, the colonies were fixed with 10% formaldehyde (VWR International) and stained with 0.05% crystal violet (Sigma-Aldrich). Colonies containing >50 cells were counted using a stereo microscope (Zeiss). The plating efficacy (PE) at 0 Gy and surviving fraction (SF) were calculated as described previously (Cojoc et al., 2015; Peitzsch et al., 2016).

### Sphere forming assay

Cells with or without treatment were plated as single cell suspension at a density of 2000-5000 cells/well depending on the cell line in 24-well ultra-low attachment plates in Mammary Epithelial Cell Growth Medium (MEBM) medium (Lonza, Germany) supplemented with 4 μg/ml insulin (Sigma-Aldrich), B27 in a dilution 1:50 (Invitrogen), 20 ng/ml epidermal growth factor (EGF) (Peprotech), 20 ng/ml basic fibroblast growth factor (FGF) (Peprotech). Spheres were analyzed 14 days after cell plating. Cell clusters were disaggregated by pipetting before analysis. Plates were automatically scanned using the Celigo S Imaging Cell Cytometer (Nexcelom). The number and size of spheres were analyzed using ImageJ 1.8.0 software. Complementary cumulative distribution function was used for the analysis of number and size of tumor spheres. Cell aggregates were discriminated from spheres based on their shape, size and structure and excluded from analysis.

### RNA isolation, cDNA synthesis, and RT-qPCR

RNA from PCa cells was isolated by RNeasy Mini kit Plus (Qiagen) according to the manufacturer’s recommendations. Briefly, cells were lysed with 350 μl of RLT buffer supplemented with 1M dithiothreitol (DTT) (Sigma) to a well of a 6-well plate that was previously rinsed with PBS. The lysate is then passed through a gDNA Eliminator spin column, which allows efficient removal of genomic DNA. 70% Ethanol was added to the flow-through to provide appropriate binding conditions for RNA, and the sample is then applied to an RNeasy spin column, where total RNA binds to the membrane and contaminants are efficiently washed away. High-quality RNA was then eluted in 70 μl of RNase-free water. The concentration of isolated RNA was measured using the spectrophotometer Nano Drop-1000 under control of 260/280 and 260/230 ratio to exclude contamination by residual phenol, guanidine, or other reagent used in the extraction protocol. Reverse transcription was done using the PrimeScript™ RT reagent Kit (Takara) according to the manufacturer’s recommendation. The volume of RNA for reverse transcription was adjusted in all samples to obtain a unified RNA concentration of at least 500 ng per sample. As technical control minus reverse transcriptase control (-RT) was used, which involved carrying out the reverse transcription step of a qRT-PCR experiment in the absence of reverse transcriptase. Quantitative real-time polymerase chain reaction (qRT-PCR) was carried out using the TB GreenTM Premix Ex TaqTM II (Takara Bio Inc., Kusatsu, Japan) according to the manufacturer’s protocol for a total reaction volume of 20 μl. Each cDNA was diluted to a working concentration of at least 15 ng/μl in RNase-free water prior to the PCR run. The qPCR cycling program was set on a StepOnePlus system (Applied Biosystems): 95°C for 10 min, 40 cycles: 95°C for 15 s, 60°C for 60 s, 72°C for 60 s followed by a melt curve to 95°C in steps of 0.3°C. All experiments were conducted using at least three technical replicates and the expression of ACTB, RPLP0, and B2M mRNA was used as reference to internal control for data normalization depending on experiment. The primers used in the study are listed in Supplementary table 2.

### Aldefluor™ assay and flow cytometry

Aldehyde dehydrogenase activity was analyzed using the Aldefluor™ assay, according to the manufacturer’s protocol (Stem Cell Technologies). In brief, cells were detached using Accutase (PAA), washed with PBS and resuspended in Aldefluor buffer. Cells were incubated with the specific ALDH inhibitor diethylaminobenzaldehyde (DEAB) in concentration 1:50 which served as a negative control. Both control and positive samples were then stained with Aldefluor reagent at the concentration 1:200 and incubated at 37°C for 30 minutes. Dead cells were excluded by 1μg/ml PI staining and doublets were excluded using the FSC-W and SSC-W function of the BD FACSDiva™ 8.0.1 software. Stained cells were excited with a blue laser (488 nm) and the analysis was performed using FITC channel. Samples were analyzed with the BD Celesta flow cytometer. A minimum of 100.000 viable cell events were collected per sample. Data were analyzed using FlowJo 10.7 software and gates were set according to the DEAB control.

### Chemical treatment

Enzalutamide, XAV939 and Zolendronic acid (Zol) were purchased from Cayman Chemical Company. DMSO was used as a drug solvent for Enzalutamide and XAV939, and corresponding concentrations of DMSO were used as controls in all the experiments, which included cell treatment. PBS was used as a drug solvent for Zol and as a control. The cells were serum starved in DMEM or RPMI with 3% FBS for 24 hours followed by treatment with XAV939 antagonist at concentration 10, 50 and 100 μM/L and with Enzalutamide inhibitor at concentration 5, 10 and 20 μM/L for 48 hours. For Zol treatment experiments the drug was used in concentration 50 and 100 μM/L for 48 hours. After 48 h in the incubator, cells were used for RNA or protein isolation. In total, at least three independent biological repeats were performed with cells at different passages.

### PC3 xenograft tumor cell recovery and subline generation

To establish xenograft tumor growth, 1×32^6^ PC-3 Luc2/RGB cells (established as previously described in (Hoffmann et al., 2020) were injected subcutaneously in 8 to 10 weeks old male immunodeficient NSG mice (NOD.Cg-Prkdcscid Il2rgtm1Wjl/SzJ; Jax, Stock 005557). The animal experiments were approved by the local animal experiment approval committee (Behörde für Justiz und Verbraucherschutz - Lebensmittelsicherheit und Veterinärwesen, assigned project No. G 80/16 and G105/16). To investigate the outgrowth capacity of spontaneous metastases after prolonged growth periods, the primary tumors were be surgically removed before they reached a volume of 1 cm^3^ or ulcerated the mouse skin. The mice were sacrificed when the relapsing tumor reached a volume of ~1 cm^3^. At necropsy, tissue of the relapsing tumor and the lung was ground in culture medium and filtered with a 100 μm strainer. The bones of the hind limbs were cut transversally in the middle of the diaphysis and the bone marrow was harvested by centrifuging for 30 sec on 5500 g (after placing the bones with the opened sides down in PCR tubes placed in 1.5 ml Eppendorf tubes). The bone marrow was resuspended in culture medium. The cell suspensions of different tissues were cultured in RPMI-1640 medium supplemented with 10% FBS, 100 U/ml penicillin and 100 μg/ml streptomycin and 1 μg/ml puromycin. Expansion of adherent tumor cell cultures was monitored in a light microscope and sub-cultivation of xenograft tumor, lung metastasis as well as bone marrow metastasis sublines was conducted as appropriate.

### Syngeneic immunocompetent tumor model

Intracardiac injection of murine prostate cancer cells RM1(BM) with or without the knockdown of *Aldh1a1* and *Aldh1a3* gene followed with immunofluorescence analysis was used to examine the ability of the cells to home to bone. Anesthetized animals were placed in a supine position for tumor cell injection. Male C57BL/6 mice (8-9 weeks old) were injected into the left ventricle with shNS, shAldh1a1, or shAldh1a3 cells (1×32^5^ cells per mouse). On day 3 post injections mice were killed by cervical dislocation, the legs from the mice were removed and knees detached from femur and tibia. Than the bones were fixed in 4% PFA overnight at 4°C. The next morning bones were washed in PBS and dried. Decalcification was performed with 2 ml of Osteosoft reagent (Merck Millipore) at 37°C for 5 days. Afterwards, the bones were embedded in OCT Tissue-Tek specimen matrix compound (Sakura) and cut for 10 μm thick sections with Cryotome. The bones were stored at −20°C. Experiments were approved by the Landesdirektion Sachsen.

### Immunofluorescence of bone metastasis on frozen sections

First, frozen sections were left at room temperature for at least 30 min and area of section was marked with Pap-Pen (Thermo Fisher Scientific). After 5 min fixation of sections with 4% PFA, they were washed and permeabilised with Triton-X-100 0.01% in PBS. After 3 times washes with PBS, the sections were blocked with Protein block Serum free DAKO (Agilent) for 30 to 45 min at RT. Incubation with primary antibodies was done in DAKO Antibody diluent with background reducing components (Agilent) at 4°C overnight. Next day, after 3 times washes with PBS, the sections were incubated with secondary antibody in DAKO Antibody diluent for 1.5 hours at RT. Afterwards, sections were washed 3 times with PBS and stained with DAPI (1 mg/ml, 1:5000 in 1xPBS) for 5 min at room temperature. Later, the slides were mounted with fluorescent mounting medium DAKO (Agilent) and left at +4°C overnight. The imaging was performed with WF Slide scanner Axioscan (Zeiss). The antibodies used in the study are represented in Supplementary table 2.

### Analysis of the TCGA patient cohort data

The publicly available TCGA PRAD dataset was downloaded from cBioportal https://www.cbioportal.org/ and analysed using SUMO software https://angiogenesis.dkfz.de/oncoexpress/software/sumo/. For evaluation of correlation with *ALDH1A1* and *ALDH1A3* the RT2 profiler PCR Array gene sets (Qiagen) were used, the Pearson correlation coefficients for all genes included in each gene set were determined, and the median correlation, coefficients were calculated. For Kaplan-Meier survival analysis, the biochemical recurrence-free survival time was determined based on provided “Days to PSA” and “Days to biochemical recurrence first” data, and the patient groups were defined by the optimal cut-off scan procedure.

### Statistical analysis

The cell survival curves were analyzed using the Statistical Package for the Social Sciences (SPSS) v23 software as described previously (Franken et al., 2006) by linear– quadratic formula S(D)/S(0) = exp−(αD + βD2) using stratified linear regression. Significance was determined by GraphPad Prism software V6. A significant difference between two conditions was recorded for *p<0.05; **p<0.01; ***p<0.001. Correlation of gene expression levels was evaluated by SUMO software using Pearson correlation coefficient. For *in vivo* mouse experiment outliers were removed by iterative Grubbs’ method with α = 0.05. p<0.05 was considered as significant by the Mann-Whitney U test.

## Results

### ALDH1A1 and ALDH1A3 regulate CSC phenotype

Previous studies have shown that ALDH enzymatic activity can be used to identify and isolate prostate cancer cells with stem cell properties (Cojoc et al., 2015). We compared the expression of seven ALDH-specific isoform genes, shown to be highly expressed in prostate cancer clinical samples, by gene expression profiling of ALDH^+^ and ALDH^−^ cell populations isolated by FACS from DU145 cells (Cojoc et al., 2015; Magnen et al., 2013). We revealed that *ALDH1A3* was significantly upregulated in the ALDH^+^ cell population. The other genes showed either minor or inconsistent differential expression in ALDH^+^ as opposed to ALDH^−^ cells (Fig. 1A). Although *ALDH1A1* showed inconsistent results in our study, it is the only isoform whose gene expression correlated with Aldefluor activity in the patient’s tissue specimens (Magnen et al., 2013). However, when we correlated the expression of *ALDH1A1* and *ALDH1A3* genes to the Aldefluor activity in four prostate cancer cell lines, ALDH1A3 showed high correlation with the fraction of ALDH^+^ cells (R^2^=0.968), ALDH1A1 did not show any (Fig. 1B). To evaluate contribution of ALDH1A1 and ALDH1A3 to the CSCs phenotype, we performed siRNA-mediated knockdown of these genes and measured Aldefluor activity by flow cytometry. This experiment confirmed that both genes equally contribute to ALDH enzymatic activity (Fig. 1C). These results suggest that regulation of the ALDH activity cannot be solely explained by the level of gene expression but also attributed to the posttranslational protein modifications as showed previously (Ibrahim et al., 2018). Since ALDH^+^ cells exhibit stem-like properties, we next analyzed an association of *ALDH1A1* and *ALDH1A3* with CSC phenotype under serum-free sphere-forming conditions. Cells with genetically silenced *ALDH1A1* and *ALDH1A3* expression showed significant decrease in spheres number and size (Fig. 2A and B). These experiments confirmed ALDH1A1 and ALDH1A3 as the essential contributors to ALDH activity and functional regulators of CSC phenotype.

**Figure 1.**
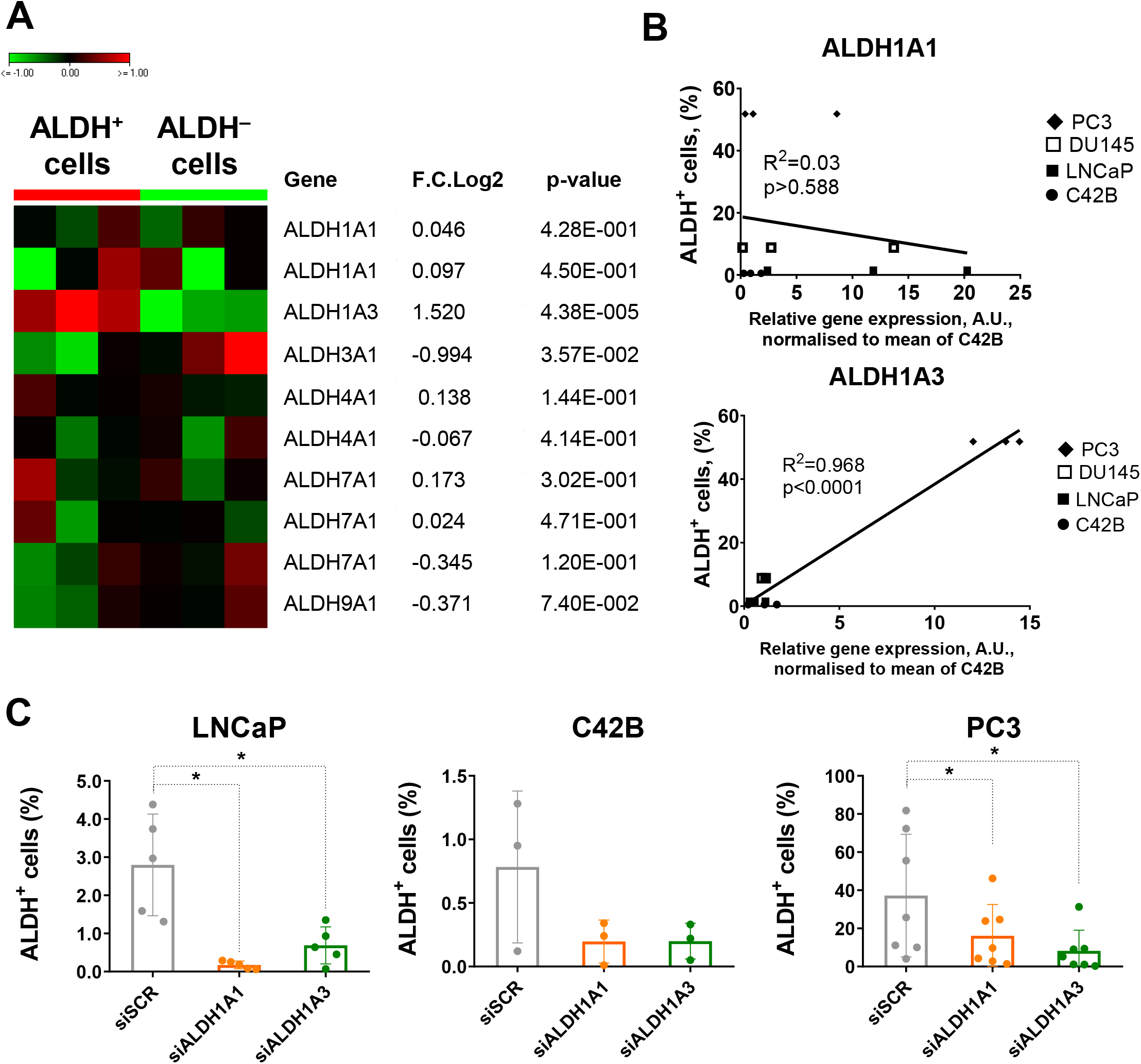
ALDH1A1 and ALDH1A3 contribute to Aldefluor activity. **A,** Expression of clinically relevant ALDH genes in ALDH^+^ and ALDH^−^ populations of DU145 cells. Significance between ALDH^+^ and ALDH^−^ samples was determined by two-tailed paired t-test. **B,** Correlation of ALDH1A1 and ALDH1A3 expression across four prostate cancer cell lines with mean size of ALDH^+^ proportion. **C,** Flow cytometry analysis of ALDH^+^ populations upon ALDH1A1 and ALDH1A3 knockdown shows a decrease of Aldefluor enzymatic activity. N≥3; Error bars = SD; Significance was determined by two-tailed paired t-test; *p<0.05.

**Figure 2.**
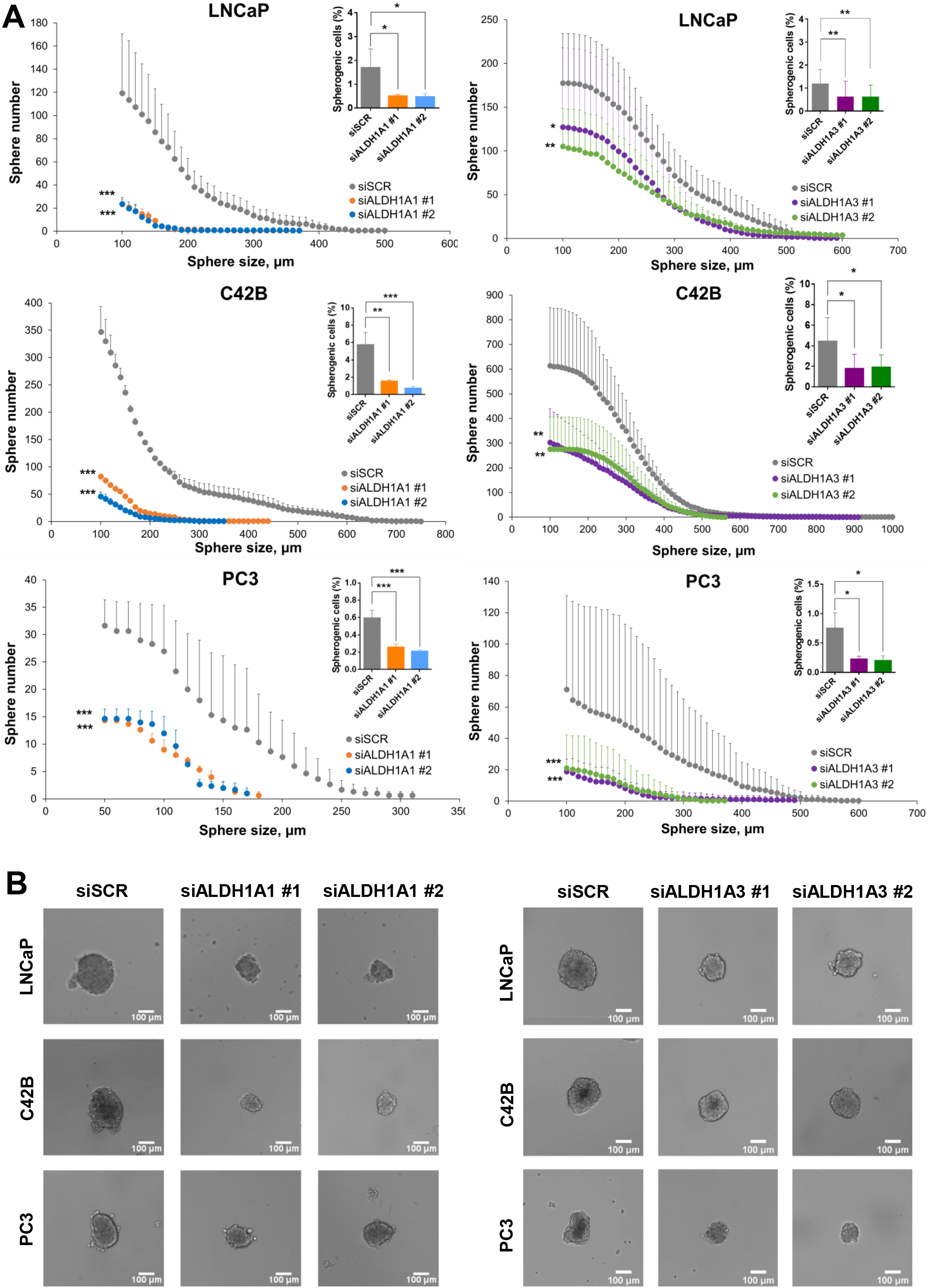
ALDH1A1 and ALDH1A3 as regulators of CSC phenotype. **A,** Complementary cumulative distribution for the number and size of tumor spheres after ALDH1A1 and ALDH1A3 depletion. The graph represents the number of spheres larger than thresholds. N≥3; Error bars = SEM; *p<0.05; **p<0.01; ***p<0.001. Bar graph represents the total number of formed tumor spheres upon ALDH1A1 and ALDH1A3 knockdown. N≥3; Error bars = SD; Significance was determined by two-tailed paired t-test; *p<0.05; **p<0.01; ***p<0.001. **B,** Representative images of prostatospheres are shown for cells transfected with scrambled siRNA (control) and with ALDH1A1 or ALDH1A3 siRNAs. Scale bar = 100μm.

### Knockdown of ALDH1A1 and ALDH1A3 results in PCa radiosensitization

One of the main features of PCa stem cells is resistance to conventional therapies, including radiation therapy (RT). Previous studies showed that ALDH^+^ prostate CSCs exhibited higher radioresistance than non-CSC counterparts (Cojoc et al., 2015). Therefore, we hypothesized that ALDH1A1 and ALDH1A3 might be involved in the RT response of PCa cells. To answer this question, we first measured our target genes’ expression in radiation resistant (RR) sublines. Those cell lines were established by applying multiple fractions of 4 Gy X-rays to the established PCa cells until a total dose of more than 56 Gy was reached as described previously (Cojoc et al., 2015). This analysis revealed that *ALDH1A1* was highly increased, whereas *ALDH1A3* was significantly downregulated in RR cells (Fig. 3A). Next, we used clonogenic radiobiological cell survival assay to analyze the relative radioresistance of cells upon siRNA-mediated knockdown of *ALDH1A1* and *ALDH1A3*. The results of these experiments demonstrated that genetic silencing of both genes with two specific siRNAs led to significant radiosensitization of PCa cells *in vitro*, with a more pronounced effect after *ALDH1A3* depletion (Fig. 3B; Supplementary Fig S1). These data indicate that both ALDH1A1 and ALDH1A3 contribute to the regulation of cell radiosensitivity.

**Figure 3.**
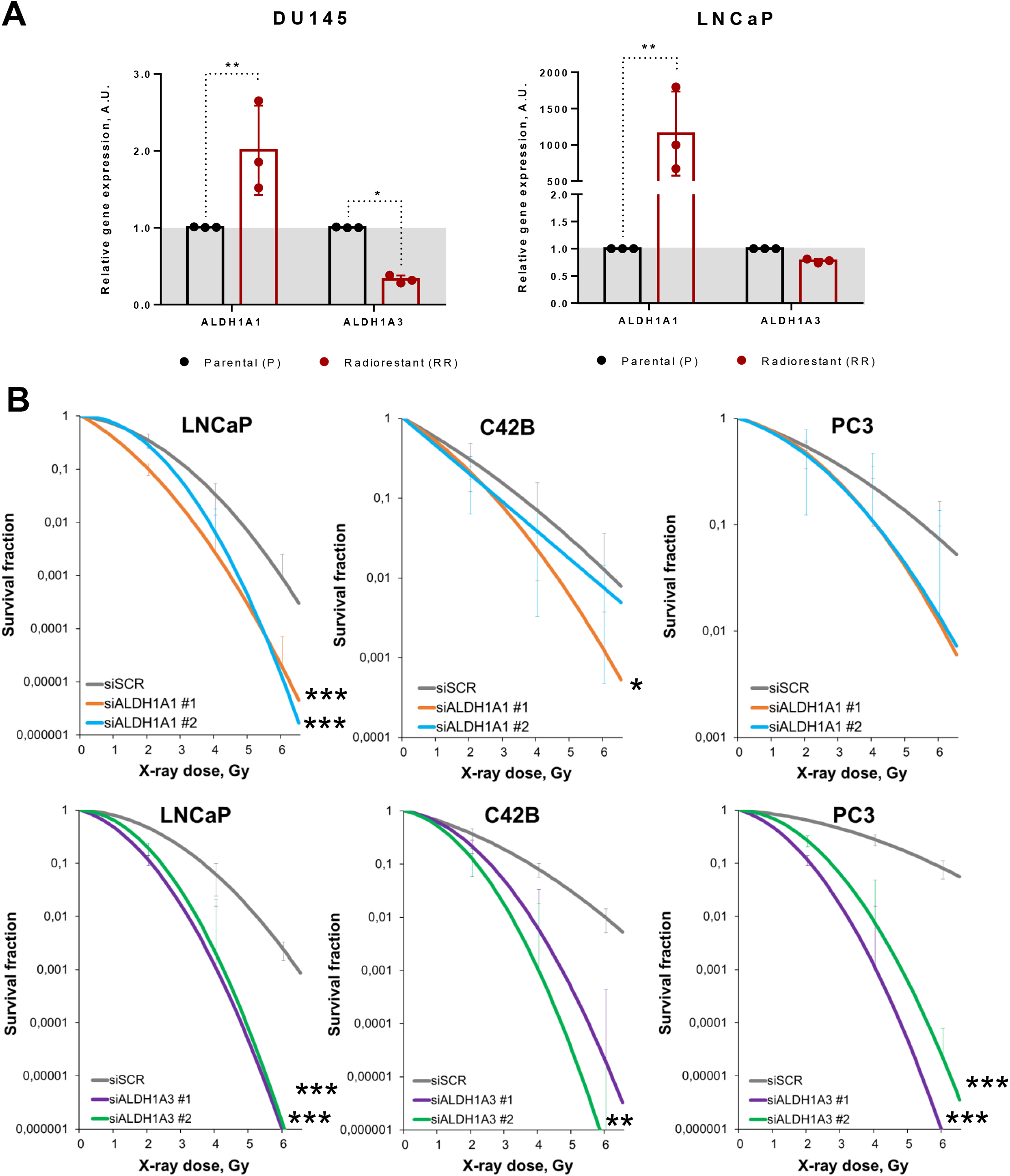
ALDH1A1 and ALDH1A3 as regulators of radioresistance. **A,** qPCR analysis of ALDH1A1 and ALDH1A3 genes expression in parental (P) and radioresistant (RR) cells; Normalized to housekeeping gene ACTB and plotted relative to parental cells; N=3; Error bars = SD. Significance was determined by two-tailed paired t-test; *p<0.05; **p<0.01. **B,** Surviving fraction of cells upon ALDH1A1 and ALDH1A3 downregulation. N= 3; Error bars = SD. Plotted lines were fitted using the linear quadratic model. Significance was calculated in SPSS software. *p<0.05; **p<0.01; ***p<0.001.

### Expression levels of ALDH1A1 and ALDH1A3 genes are mutually regulated

Despite ALDH1A1 and ALDH1A3 have a similar physiological function and both appear as regulators of CSC properties and radioresistance (Vassalli, 2019), we wondered if one of the isoforms is more essential for PCa development and progression than the other. We first determined the correlation of those genes to each other in the publicly available provisional PCa cohort (PRAD) from The Cancer Genome Atlas (TCGA) (n=490). The analysis revealed a weak negative correlation between those genes (*r =* −0.19485, p<0.05) (Fig 4A). This observation motivated us to investigate the relationship between *ALDH1A1* and *ALDH1A3* further in knockdown experiments. Considering that the two siRNAs used for previous experiments showed similar trends, we used the pooled siRNA for further investigations and found that genetic silencing of *ALDH1A1* led to downregulation of *ALDH1A3*; however, the depletion of *ALDH1A3* increased *ALDH1A1* mRNA expression in all three PCa cell lines (Fig. 4B).

**Figure 4.**
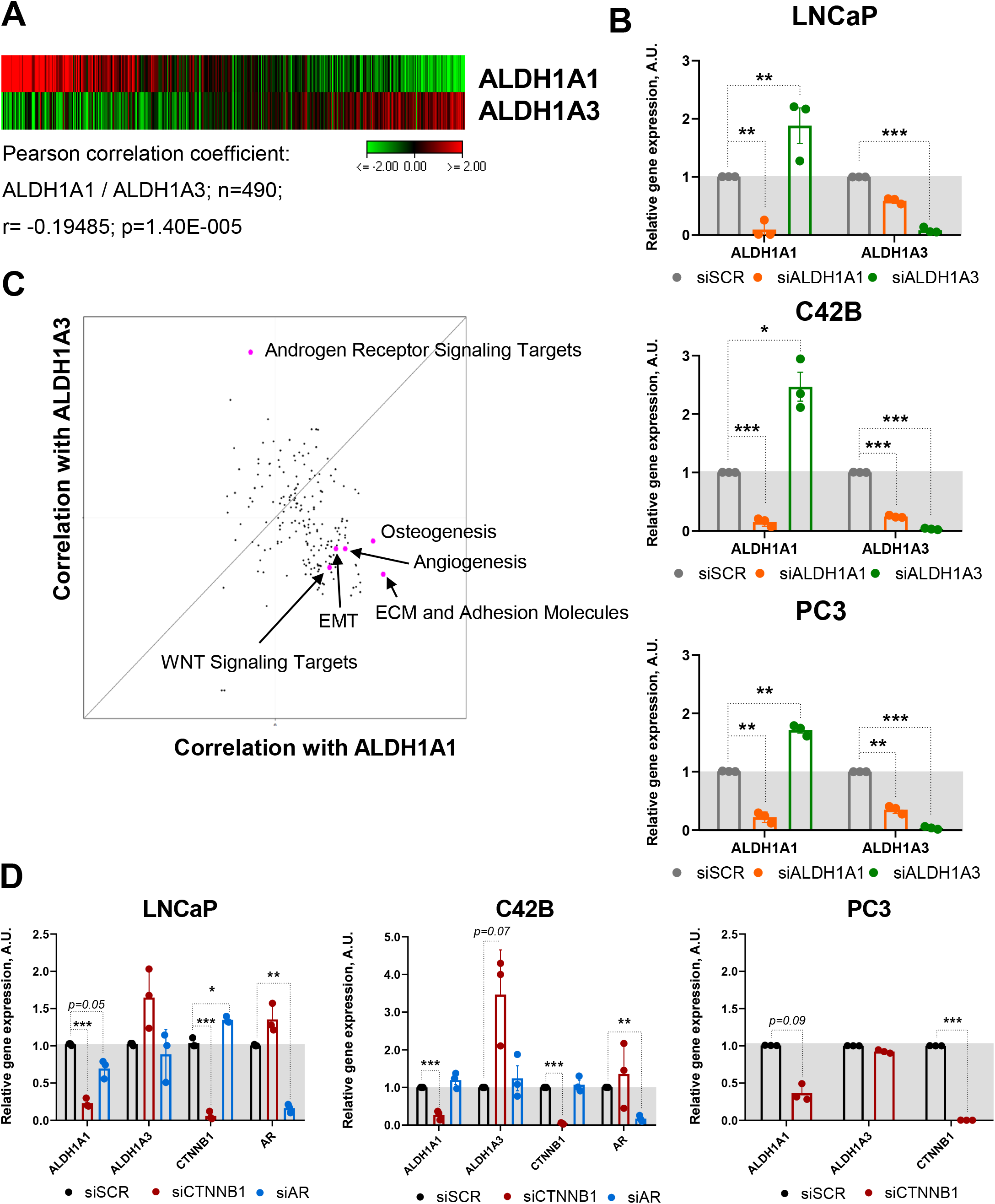
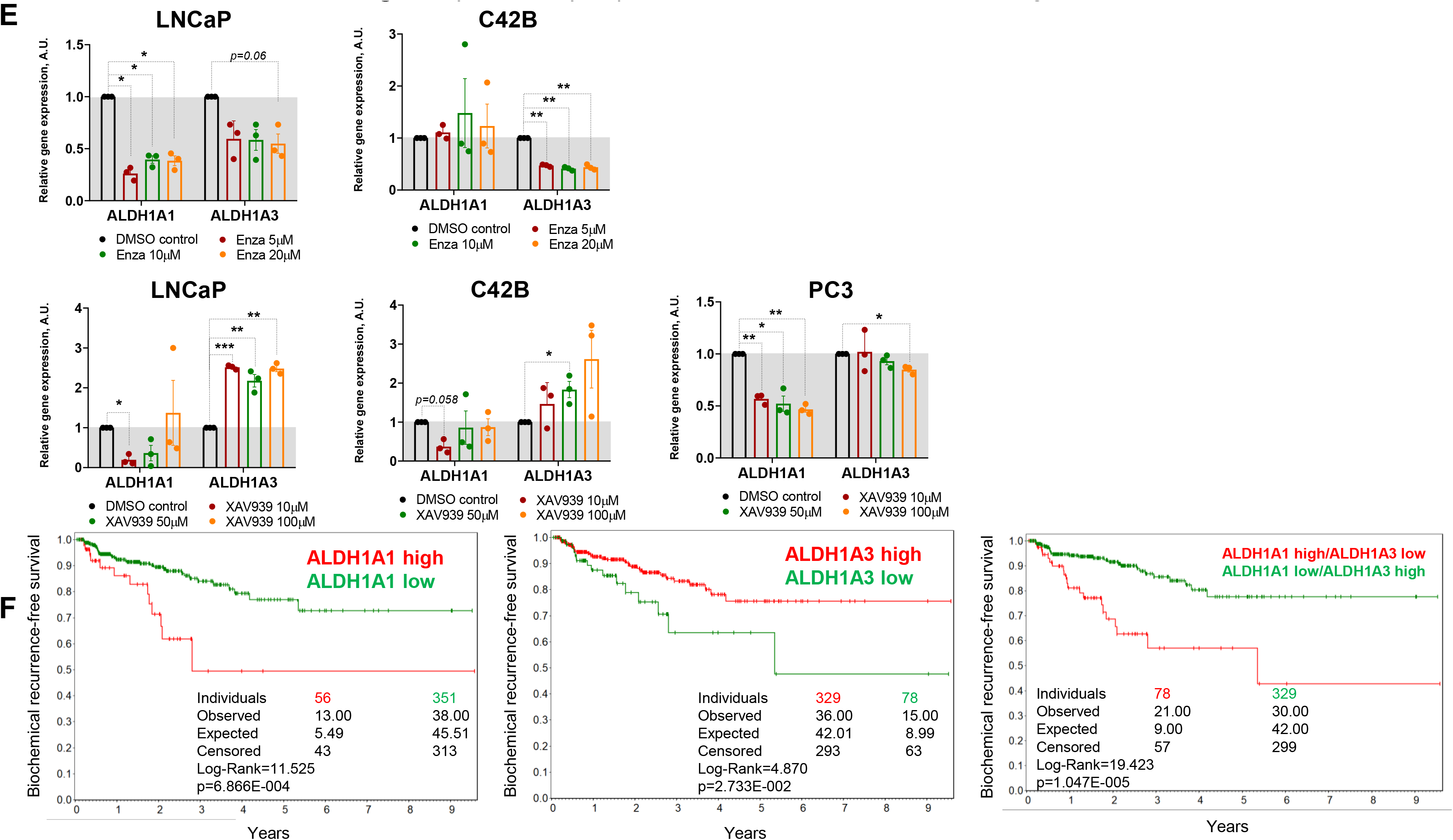
Expression of the ALDH1A1 and ALDH1A3 genes is interconnected. **A,** Analysis of the TCGA dataset for patients with PCa (n=490) showed a weak negative correlation of ALDH1A1 and ALDH1A3 genes. Significance was determined using the Pearson coefficient. **B,** qPCR analysis of ALDH1A1 and ALDH1A3 expression after reciprocal knockdown and scrambled control. Normalized to housekeeping gene ACTB and plotted relative to the control siSCR sample. Significance calculated by two-tailed paired t-test. N= 3; Error bars = SD. **p<0.01; ***p<0.001. **C,** The correlation of the common molecular pathways with ALDH1A1 and ALDH1A3 in a provisional prostate cancer TCGA dataset (n=490). **D,** Relative mRNA expression of ALDH1A1, ALDH1A3, CTNNB1 and AR upon the knockdown of CTNNB1 and AR genes. Normalized to housekeeping gene RPLP0 and plotted relative to the control siSCR sample. Significance calculated by two-tailed paired t-test. N= 3; Error bars = SD. **p<0.01; ***p<0.001. Expression of the ALDH1A1 and ALDH1A3 genes is interconnected. **E,** Analysis of ALDH1A1 and ALDH1A3 genes expression upon inhibition of WNT signaling pathway with XAV939 inhibitor and AR signalling with Enzalutamide. Normalized to housekeeping gene RPLP0 and plotted relative to the DMSO control sample. The cells were serum starved in DMEM or RPMI medium with 3% FBS for 24 hours followed by treatment with XAV939 or Enzalutamide antagonist at different concentrations. Significance calculated by two-tailed paired t-test. N= 3; Error bars = SD. *p<0.05; **p<0.01; ***p<0.001. **F,** The Kaplan-Meier analyses of BRGC for TCGA PRAD patients stratified by the most significant cutoff for ALDH1A1 and ALDH1A3 expression levels.

The growing body of evidence proposes that ALDH1A1 and ALDH1A3 both functionally contribute to various steps in the tumor development and metastatic process (Rodriguez-Torres and Allan, 2016); nevertheless, they might function in different ways. Our previous studies showed that activation of the WNT/β-catenin signaling pathway increased β-catenin/TCF4-dependent transcription of the *ALDH1A1* gene associated with gaining of EMT features and migratory behavior (Cojoc et al., 2015). To further explore the role of ALDH genes in prostate cancer, we analysed the association of the common molecular pathways with ALDH1A1 and ALDH1A3 in the TCGA gene expression dataset. On one hand, our analysis revealed a positive correlation of the *ALDH1A1* gene with several gene sets related to cancer progression, e.g. WNT signalling, angio- and osteogenesis, extracellular matrix and adhesion molecules. On the other hand, ALDH1A3 was strongly associated with AR signalling (Fig. 4C; Supplementary Fig S2) that can be potentially attributed to the previously reported regulation of *ALDH1A3* expression by *AR* (Trasino et al., 2007). To evaluate the contribution of these genes in the regulation of cancer progression, we profiled cells with *ALDH1A1* and *ALDH1A3* knockdown by an extracellular matrix and adhesion RT² Profiler PCR Array covering 84 genes important for cell-cell and cell-matrix interactions. The experiment showed that most of the genes were similarly regulated in both knockdown conditions (Supplementary Fig S3). To investigate the link between ALDH genes and either WNT or AR signaling, we genetically silenced *AR* and *CTNNB1* genes and measured the mRNA expression level of *ALDH1A1* and *ALDH1A3*. These experiments showed that upon *AR* downregulation, the expression of *ALDH1A1* and *ALDH1A3* did not significantly change in two prostate cancer cell models (Fig. 4D). Downregulation of the *CTNNB1* gene led to the significant decrease of *ALDH1A1* expression in all cell lines with a smaller fold change in PC3 cells that was already demonstrated previously (Cojoc et al., 2015). Contrary, *ALDH1A3* was upregulated in the *CTNNB1*-depleted condition except in PC3 cells (Fig. 4D).

We also looked for a drugable approach to achieve ALDH genes downregulation. Therefore, we validated our findings by using the clinically approved antiandrogen – Enzalutamide. We also used the WNT pathway inhibitor, XAV939, showed to be efficient for inhibition of β-catenin (Cojoc et al., 2015; Peitzsch et al., 2016). Those experiments confirmed the results obtained with genetic silencing (Fig. 4E). To investigate the predictive value of ALDH1A1 and ALDH1A3, we analysed biochemical recurrence-free survival of TCGA PRAD patients stratified based on the expression of those two genes. This analysis demonstrated that *ALDH1A1* and *ALDH1A3* gene expression results in an opposing predictive value for biochemical recurrence-free survival (BRFS). The same trend was observed for disease-free survival (DFS) (Supplementary Fig S4). Combining ALDH1A1-high with ALDH1A3-low signatures results in a higher predictive value for BRFS (Fig. 4F).

### ALDH genes differentially regulate prostate cancer metastasis

As previously noted, ALDH1A1 and ALDH1A3 showed an opposite correlation with the clinical outcome; hence, we wondered how these genes contribute to tumor progression. Metastasis to the bone is the leading cause of death for prostate cancer patients. To colonize a metastatic site, tumor cells have to detach from the primary tumor, intravasate, survive in the circulation, and finally, extravasate and invade the target tissue. To investigate the role of ALDH1A1 and ALDH1A3 for tumor cell survival in the blood stream and during the extravasation process *in vivo*, we implemented the Zebrafish (*Danio rerio)* model to xenograft with human prostate cells. Zebrafish represent a powerful tool for cancer research: human and zebrafish genomes share gene orthology for 82% of disease-causing genes (Amawi et al., 2021; Howe et al., 2013; Letrado et al., 2018; Terriente and Pujades, 2013), and transparent embryos can be injected with cancer cells at various locations, allowing the live observation of tumor cells (Dietrich et al., 2021). In our study, we have used the embryonic and larval in *vivo* model to analyze the role of *ALDH1A1* and *ALDH1A3* genes in human cancer dissemination.

First, we established two color-coded PC3 cell lines with gene vectors encoding for the red fluorescent protein tdTomato or the green fluorescent protein GFP. Next, we validated that both fluorescent proteins do not affect tumor cell extravasation. To do so, we intravenously co-injected PC3-GFP and PC3-tdTomato cells into the DoC of the *Tg(kdrl:CFP)* endothelial reporter transgenic Zebrafish at 2dpf. To visualize vital and extravasated cells at 3dpi, we performed high-resolution imaging of the whole tail region including the caudal hematopoietic tissue (CHT), the site of blood formation at this developmental stage, using a Dragonfly Spinning disk confocal microscope. We quantified the number of survived cells in the bloodstream and of extravasated cells in the tail region after confirming no significant specific effect of GFP or tdTomato expression on cell survival and extravasation (Supplementary Fig S5A).

Next, we performed the siRNA-mediated knockdown of *ALDH1A1* or *ALDH1A3* in PC3-tdTomato cells. PC3-GFP cells transfected with scrambled siRNA (siSCR) were used as control. Then, we intravenously co-injected siSCR PC3-GFP and siALDH1A1 or siALDH1A3 PC3-tdTomato cells into the DoC of the *Tg(kdrl:CFP) Z*ebrafish larvae at 2 dpf. The survived and extravasated cells were analyzed at 3dpi as described above. The data showed that cells depleted for *ALDH1A1* had a survival disadvantage in the blood flow opposing to *ALDH1A3* knockdown cells. We evaluated the extravasation potential of PC3 cells with or without *ALDH1A1* and *ALDH1A3* depletion by counting the number of extravasated cells in the tail region. Cells with suppressed *ALDH1A3* expression showed higher extravasation capacities than the siSCR control. Cells with *ALDH1A1* knockdown did not show any differences in extravasation capacities. Moreover, we performed the same evaluation for tumor cells upon knockdown of β-catenin/CTNNB1, which has been shown to positively regulate *ALDH1A1* expression (Cojoc et al., 2015; Gorodetska et al., 2019; Peitzsch et al., 2016) (Fig. 4D and Supplementary Fig S5B) and negatively *ALDH1A3* expression (Fig. 4D). These experiments revealed decreased survival and extravasation of cells upon *CTNNB1* depletion (Fig. 5A and B).

**Figure 5.**
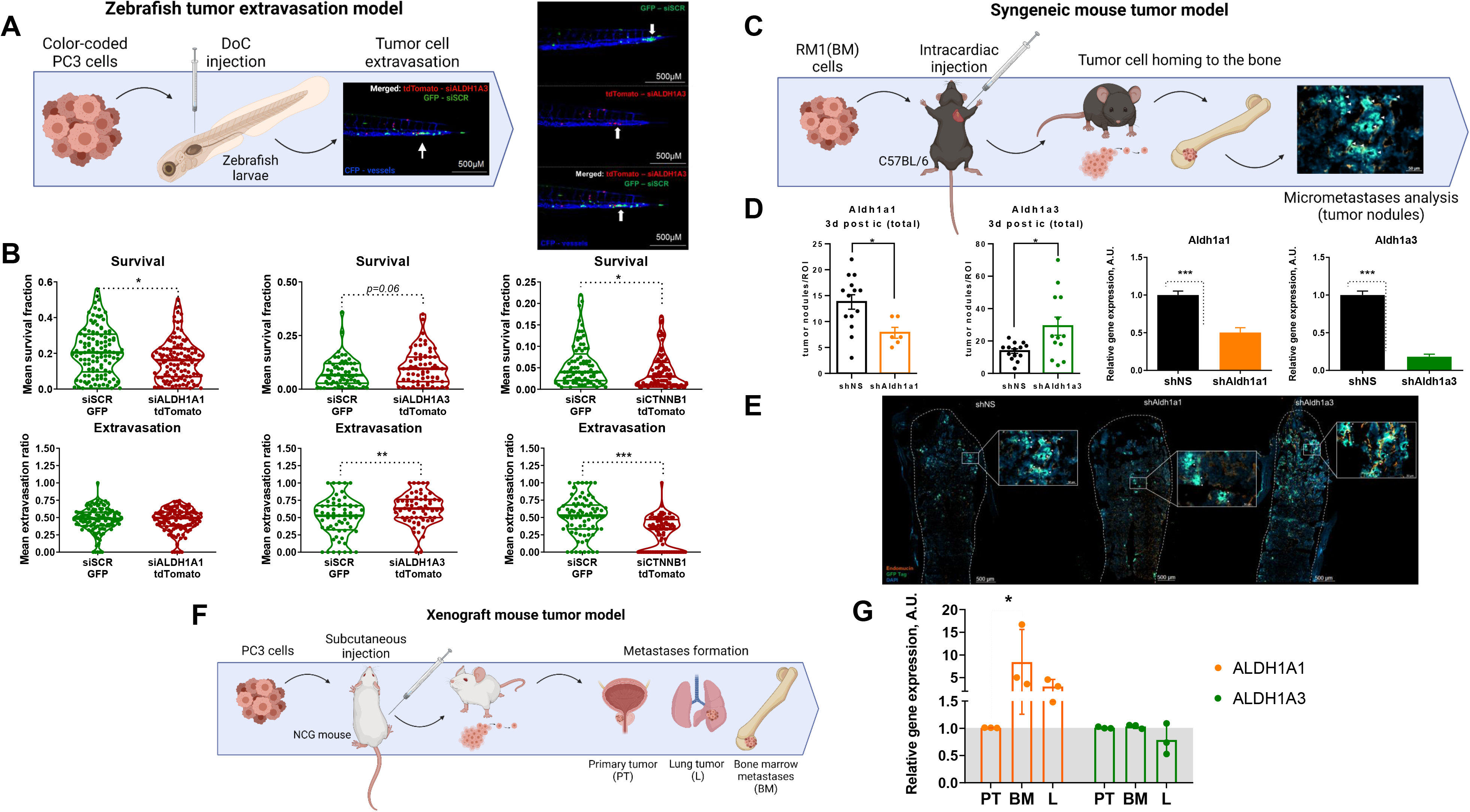
ALDH genes differentially regulate metastasis formation. **A,** Schemata of the experiment for the Zebrafish tumor extravasation model. Representative fluorescent images of the zebrafish tail. CFP – vessels; tdTomato – color-coded prostate cancer PC3 cells transfected with ALDH1A3 siRNA; GFP – color-coded prostate cancer PC3 cells transfected with siSCR. Scale bars = 500μM. **B,** The survival and extravasation potential of color-coded PC3 cells upon ALDH1A1, ALDH1A3 and CTNNB1 knockdown in the tail region. Data represents the mean of three biological repeats pooled together. ALDH1A1 experiment: N=116 fish (siSCR), 114 fish (siALDH1A1); ALDH1A3 experiment: N=61 fish (siSCR), N=63 fish (siALDH1A3); CTNNB1 experiment: N=77 fish (siSCR), N=81 fish (siCTNNB1). Statistics were performed using two-sided Mann–Whitney U test. *p<0.05; **p<0.01; ***p<0.001. **C,** Schemata of the experiment for the syngeneic mouse tumor model. After intracardiac injection of mouse prostate cancer RM1(BM) cells with or without of Aldh1a1 and Aldh1a3 depletion. Three days post intracardiac injections the formation of tumor nodules can be determined in the bones by immunofluorescence analysis. **D,** The number of tumor nodules formed in the bone tissue upon knockdown of Aldh1a1 or Aldh1a3 in the syngeneic immunocompetent mice was analysed by immunofluorescence (N = 4 mice/group; N of analysed bone slides: shNS = 14, shAldh1a1 = 6, shAldh1a3 =13; ROI (region of interest) = one bone slide). Data represents the mean of 6-14 analysed slides for one group. Outliers were removed by iterative Grubbs’ method with α = 0.05. Statistics were performed using two-sided Mann–Whitney U test. Error bars = SEM; *p<0.05. Validation of knockdown efficacy in the mouse prostate cancer cell line RM1(BM) transfected with shRNA against Aldh1a1 and Aldh1a3. Normalized to housekeeping gene Gapdh and plotted relative to the control shNS sample. Error bars = SD. Significance was determined by two-tailed paired t-test; ***p<0.001. **E,** Representative immunofluorescence images of the formed tumor modules in the shAldh1a1, shAldh1a3 and control shNS samples. Scale bar 500um and 50uM. **F,** Schemata of the experiment for the xenograft mouse tumor model. After subcutaneous engraftment of human prostate cancer PC3 cells, primary prostate tumors (PT), bone marrow (BM) and lung (L) metastasis were formed. Small pieces of surgically excised tumors were cultured in vitro and gave rise to sublines PC3-PT(derived from the primary tumor), PC3-BM (derived from BM metastasis) and PC3-L (derived from lung metastasis). **G,** qPCR analysis of ALDH1A1 and ALDH1A3 expression in the PC3 cells originating from different sites: primary tumors (PT), bone marrow (BM) and lung metastasis (L). Normalized to housekeeping gene RPLP0 and plotted relative to the PT sample. Significance calculated by two-tailed paired t-test. N= 3; Error bars = SD. *p<0.05.

We then tested whether *ALDH1A1* and *ALDH1A3* are involved in regulating tumor cells homing to the bones and bone marrow colonization *in vivo*. In this regard, we established a syngeneic murine prostate cancer model by intracardiac injection of RM1(BM) murine prostate cancer cells with bone metastases take rate over 95% into the immunocompetent C57BL/6 mice as discussed previously (Power et al., 2009). We first transfected RM1(BM)-GFP cells with plasmid vectors encoding shRNA against *Aldh1a1* or *Aldh1a3* to generate stable lines with suppressed target gene expression (shAldh1a1 or shAldh1a3). RM1(BM) cells transfected with a non-silencing shRNA (shNS) were used as control. shAldh1a1 and shAldh1a3 cells showed a reduction in expression of target genes by 50% and 80%, compared with that of the control shNS cells, analyzed by qPCR. We examined the ability of the resulting shAldh1a1 or shAldh1a3 knockdown cells to metastasize to the skeleton after being injected into the left ventricles of male C57BL/6 mice. Three days post intracardiac injections, we sacrificed the animals and isolated the hind limb (femurs and tibiae). The homing of RM1(BM) cells to the bones and their growth therein was followed by immunofluorescence microscopy analysis of GFP-positive tumor nodule formation in bone marrow tissue. Bone marrow endothelium was stained with anti-endomucin antibody. We evaluated the metastatic potential of cells upon shAldh1a1 or shAldh1a3 knockdown conditions, as well as non-silencing control, by counting the tumor nodules (Fig. 5C). This experiment showed that cells depleted for *Aldh1a1* formed lower number of tumor nodules when compared to the control. At the same time, *Aldh1a3* suppressed cells exhibited a higher number of tumor nodules formed (Fig. 5D and E).

To further investigate the role of ALDH genes in metastasis, we measured the expression of *ALDH1A1* and *ALDH1A3* genes in the PC3 cells originating from different metastatic sites (Labitzky et al., 2020). Briefly, PC3 cells were first subcutaneously injected into immunodeficient NSG mice, and small pieces of surgically excised xenograft tumors (PT), as well as spontaneous lung (L) and bone marrow (BM) metastases were used for *in vitro* propagation of sublines PC3-PT, PC3-L, and PC-BM, respectively (Fig. 5F). Our analysis revealed an overexpression of the *ALDH1A1* gene in the bone marrow metastatic cells, although the expression of *ALDH1A3* was not significantly altered. These findings revealed a high correlation of *ALDH1A1* expression with metastatic load (Fig. 5G).

### ALDH proteins differentially regulate clinical outcome

To gain insights into its potential clinical importance of our findings, we investigated the expression of ALDH1A1 and ALDH1A3 proteins in benign prostatic tissues, primary prostate cancer tissues, tissues from locally advanced or recurrent PCa, lymph node and distant metastasis. The data demonstrate that the expression of ALDH1A1 increases during PCa progression and is highest in metastases. In contrast, ALDH1A3 was highly expressed in primary and locally advanced tissue samples but not in metastasis (Fig. 6A and B). Moreover, the study confirmed that high ALDH1A1 expression on primary tumors predicts disease recurrence revealing a 5-year-progression free survival of 69.9%, 67.3% and 50% of patients harboring tumor with ALDH1A1 negative, low/moderate and high expressing tumors, respectively, while ALDH1A3 expression in primary tumors did not show a significant correlation with patients’ outcome (Fig. 6C). Comparing the initial (i) PSA serum level preoperatively between patients with tumors overexpressing ALDH1A1 and ALDH1A3, we observed a significantly reduced iPSA level in patients with ALDH1A3 overexpressing tumors (p<0.001) (Supplementary Fig S6A). There was no significant difference in the nuclear AR expression between tumor with or without ALDH1A1 or ALDH1A3 expression (Supplementary Fig S6B).

**Figure 6.**
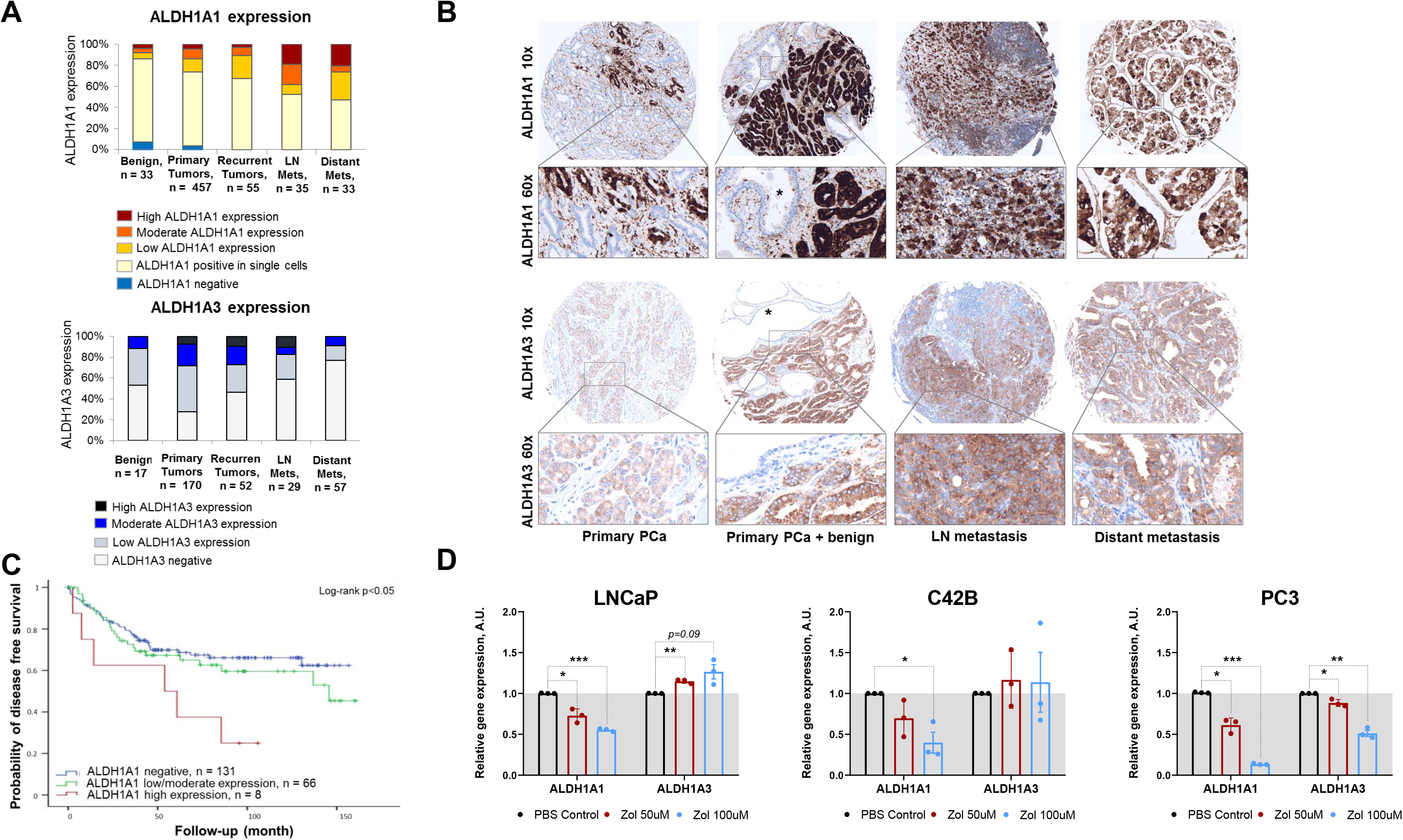
ALDH proteins as prognostic markers. **A,** The distribution of ALDH1A1 (N=613) and ALDH1A3 (N=325) expression in the benign prostatic hyperplasia (BPH), primary prostate cancer tissues, recurrent tumor, lymph node and distant metastasis cells. **B,** Representative images showing ALDH1A1 and ALDH1A3 expression in prostate cancer tissue at 10x and 60x magnification. Heterogeneous, overall low ALDH1A1 and low ALDH1A3 expression in primary PCa. Strong ALDH1A1 and ALDH1A3 expression in primary prostate cancer adjacent to normal glands lacking (*) expression. ALDH1A1 and ALDH1A3 expression in an example of lymph node and distant metastasis, respectively. **C,** The Kaplan-Meier analysis of disease-free survival of patients with high (red) compared to low/moderate (green) and undetectable (blue) ALDH1A1 expression level. N= 205; p<0.05. **D,** Relative mRNA expression of ALDH1A1 and ALDH1A3 upon Zoledronic acid (Zol) treatment. Normalized to housekeeping gene RPLP0 and plotted relative the PBS control sample. Significance calculated by two-tailed paired t-test. N= 3; Error bars = SD. *p<0.05; **p<0.01; ***p<0.001.

Bisphosphonates are a gold standard for the medical management of metastatic bone disease to inhibit bone resorption (Saad et al., 2006). Zol prevents bone complications as it inhibits osteoblastic and osteolytic metastases of prostate cancer and therefore improves life quality (Finianos and Aragon-Ching, 2019). We treated PCa cells with Zol and measured the mRNA expression of *ALDH1A1* and *ALDH1A3*. Expression of the *ALDH1A1* gene was inhibited in all analyzed cell lines in a dose-dependent manner, while *ALDH1A3* expression was upregulated in lymph-node metastatic cells (LNCaP) and downregulated in bone metastatic cells (PC3). Overall, these data suggest that Zol treatment is associated with changes in *ALDH1A1* and *ALDH1A3* expression (Fig. 6D).

### ALDH genes regulate metastasis formation through TGFB1 and MMPs

We further investigated the role of *ALDH1A1* and *ALDH1A3* in osteogenic differentiation. For this, we picked five genes from the Human Osteogenesis RT² Profiler PCR Array highly expressed in bone metastatic cell line PC3 compared to the lymph node metastasis-derived LNCaP cells (Supplementary Fig S7A). We validated by qPCR whether depletion of *ALDH1A1* and *ALDH1A3* affects these osteogenesis-related genes. Among those five genes, only the transforming growth factor-beta 1 (*TGFB1*) gene showed significant changes (Supplementary Fig S7B). TGFβ1 is one of the critical inducers of EMT, a complex program implicated in carcinogenesis and metastatic progression. Our experiment showed that *ALDH1A1* downregulation in LNCaP and C42B led to a decrease of *TGFB1* gene expression; on the other hand, *ALDH1A3* depletion increased the *TGFB1* level. Interestingly, *TGFB1* expression is regulated differently in the analyzed cell lines in response to ALDH gene knockdown. The correlation between ALDH gene depletion and *TGFB1* level is highest in androgen-sensitive LNCaP cells. It is becoming less pronounced in C42B cells, whereas *TGFB1* level in PC3 cells is not regulated in response to *ALDH1A1* and *ALDH1A3* knockdown (Fig. 7A). Moreover, *TGFB1* positively correlates with *ALDH1A1* gene and negatively correlates with *ALDH1A3* expression in the TCGA gene expression dataset for 490 PCa patients (Supplementary Fig S7C). Notably, none of the knockdown conditions altered the expression of *SMAD3*, which led us to the conclusion that regulation occurs through the non-SMAD pathway (Fig. 7A). Based on these data, the present findings demonstrate that *ALDH1A1* and *ALDH1A3* differently regulate expression of *TGFB1* gene, one of the major regulators of tumor metastatic dissemination.

**Figure 7.**
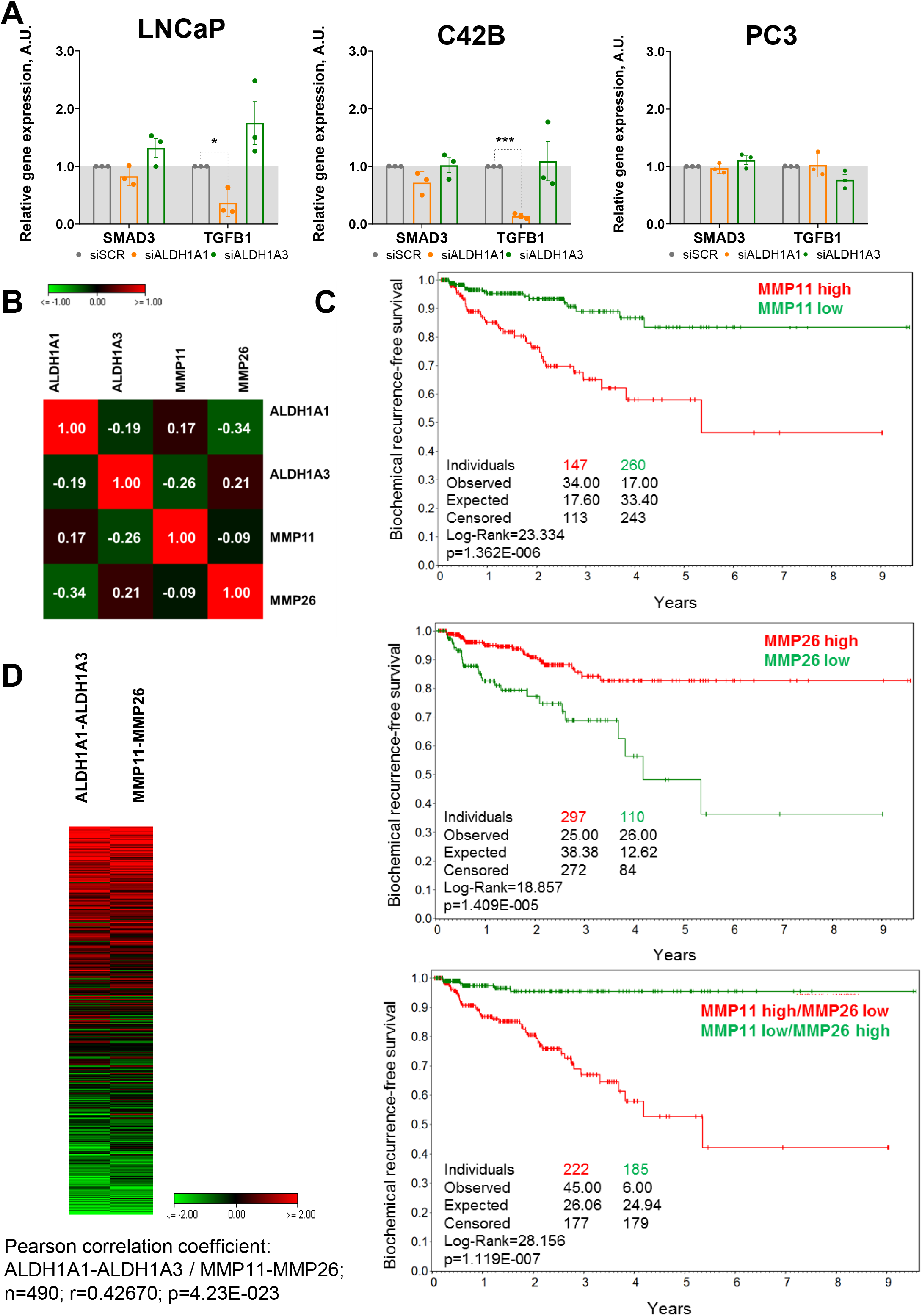
ALDH genes regulate PCa metastasis by TGFB1 and MMPs. **A,** Analysis of the expression of EMT-related genes SMAD3 and TGFB1 upon inhibition of ALDH1A1 and ALDH1A3. Normalized to housekeeping gene ACTB and plotted relative to the control siSCR sample. Significance calculated by two-tailed paired ttest. N= 3; Error bars = SD. *p<0.05; **p<0.01. **B,** Pearson correlation of ALDH1A1, ALDH1A3, MMP11 and MMP26 with each other in TCGA PRAD dataset (n=490). **C,** The Kaplan-Meier curves analysis of BRFS for TCGA PRAD patient stratification by the most significant cutoff for MMP11 and MMP26 expression level. **D,** Analysis of the TCGA dataset for patients with PCa (n=490) showed a strong positive correlation of combined ALDH1A1 high / ALDH1A3 low and MMP11 high / MMP26 low genes expression. Significance was determined using the Pearson coefficient.

Matrix metalloproteinases (MMPs), enzymes involved in the degradation of the extracellular matrix, play an important role in cancer-associated bone remodeling (Gong et al., 2014). Therefore, we closely looked at the expression of MMPs in an extracellular matrix and adhesion RT² Profiler PCR Array (Supplementary Fig S3). Several MMPs family members were substantially affected in response to ALDH1A1 and/or ALDH1A3 depletion (Supplementary Fig S8A). We analyzed all MMPs for their association with a good and bad prognosis in the PCa TCGA gene expression dataset. We identified MMP11 and MMP26 allowing the best stratification of patients (Supplementary Fig S8B). Moreover, the *MMP11* gene was also upregulated upon *ALDH1A3* expression in the qPCR array (Supplementary Fig S7A). We also found in this dataset a moderate negative correlation between *ALDH1A1* and *MMP26*, and between *ALDH1A3* and *MMP11*. In this analysis, *ALDH1A1* showed a significant positive correlation with *MMP11* and *ALDH1A3* with *MMP26* (Fig. 7B). We also performed the Kaplan–Meier analysis of biochemical recurrence-free survival of TCGA PRAD patient’s stratification by MMP11 and MMP26 expression level and found that these two genes have opposite predictive value similarly to ALDH1A1 and ALDH1A3 (Fig. 7C). Combining MMP11-high with MMP26-low gene expression leads to a higher predictive value for biochemical recurrence-free survival (Fig. 7C). Consistent with these findings, we found high positive correlation of combined ALDH1A1 high / ALDH1A3 low and MMP11 high / MMP26 low genes expression (*r =* 0.42670, p<0.05) (Fig. 7D). Based on these observations, we hypothesized that ALDH1A1 and ALDH1A3 might differentially affect metastasis formation through regulation of TGFB1 gene expression and MMP levels.

## Discussion

The CSC populations are one of the main drivers of prostate cancer progression, resistance to conventional therapies and metastases formation (Peitzsch et al., 2017). Several markers have been discovered for prostate CSCs identification, including ALDH activity, which was used for isolation of ALDH^+^ PCa cells with high clonogenic and tumorigenic capacities (Li et al., 2010). Numerous ALDH genes are contributing to ALDH enzymatic activity; however only seven of them showed clinical relevance (Magnen et al., 2013).

In our study, we found that ALDH1A1 and ALDH1A3 genes functionally regulate CSC properties and radiation sensitivity of PCa. We showed that ALDH1A1 is upregulated in bone marrow metastases of a xenograft tumor model, enhances extravasation of cancer cells in Zebrafish model and promotes the formation of tumor nodules in the bones of syngeneic immunocompetent mice model. Contrary to ALDH1A1, ALDH1A3 suppresses extravasation and survival in the blood stream of tumor cells in zebrafish and inhibits formation of tumor nodules in the mice bone marrow. We revealed a negative correlation between ALDH1A1 and ALDH1A3 expression in a publicly available PCa dataset and demonstrated that ALDH1A1 and ALDH1A3 have opposing predictive value for BRFS. We showed that treatment of PCa cells with Zol, a certified drug for the treatment of patients with progressive bone metastasis from PCa, reduced ALDH1A1 and increased ALDH1A3 expression levels. These data suggest that the previously reported inhibitory effect of Zol treatment on bone metastases (Corey et al., 2003) can be partially attributed to the regulation of ALDH1A1 and ALDH1A3 expression.

Our previous study showed high expression of ALDH1A1 in prostate tumor compared to the normal adjacent tissue (Cojoc et al., 2015). Consistently with those findings, Magnen and co-authors found an association of ALDH1A1 with higher-grade tumors (Magnen et al., 2013). A recent study by Federer-Gsponer also indicated patterns of mutual exclusivity for ALDH1A1 and ALDH1A3 expression across several datasets (Federer-Gsponer et al., 2020). However, the relationship between ALDH genes and bone metastatic tumor growth was unknown. To our knowledge, this is the first study suggesting an association of ALDH1A1 with the metastatic burden, elucidating the role of ALDH genes in the metastatic spread, and homing to the bone.

We also found that ALDH1A1 is highly correlating with metastasis-related pathways, such as WNT signalling, angio- and osteogenesis, and extracellular matrix and adhesion molecules, whereas ALDH1A3 is highly associated with AR signalling targets. Literature data show that inhibition of AR with either antiandrogen bicalutamide or siRNA decreases ALDH1A3 expression and treatment with dihydrotestosterone (DHT) increases the ALDH1A3 mRNA levels (Trasino et al., 2007). We also demonstrated that AR inhibition with Enzalutamide positively affects ALDH1A1 and ALDH1A3 gene expression, when genetic silencing of AR had no effect on ALDH1A1 and ALDH1A3 gene expression. On the other hand, depletion of β-catenin/CTNNB1 positively influences ALDH1A1 but negatively ALDH1A3 mRNA level. This opposite regulation of ALDH genes by β-catenin can be associated with their distinct role during PCa development. Previous findings showed that at the initial stages of PCa, AR signalling suppresses the transcription of β-catenin/WNT target genes, while β-catenin/WNT signalling stimulates the expression of AR targets (Pakula et al., 2017). Metastatic prostate cancers are known to become AR-negative disease associated with poor prognosis (Formaggio et al., 2021). At the same time, WNT/β-catenin signaling pathway is highly activated in AR negative PCa (Wan et al., 2012). A growing body of evidence supports the hypothesis that WNT/β-catenin signalling pathway promotes metastatic castration-resistant prostate cancer (mCRPC) by a complex interaction with AR signalling (Patel et al., 2020; Ramamurthy et al., 2018; Schweizer et al., 2008; Tsao et al., 2021). WNT/β-catenin axis is a known positive regulator of PCa metastatic dissemination (Patel et al., 2020) and treatment resistance (Yeh et al., 2019). Our previous study demonstrated that β-catenin/TCF complex directly regulates ALDH1A1 expression and promotes radiation resistance (Cojoc et al., 2015). Our present study confirmed these findings by genetic silencing and chemical inhibition of AR and CTNNB1 genes. All in all, our data indicate that ALDH1A1 and ALDH1A3 contribute to PCa progression at different stages of tumor development under AR and β-catenin-dependent regulation. ALDH genes play a diverse role in PCa development, with ALDH1A1 becoming dominant in later stages when PCa cells gain androgen independence.

We clinically validated our findings on a cohort of PCa patients. Our data support the clinical significance of ALDH1A1 and ALDH1A3, demonstrating that ALDH1A1 but not ALDH1A3 is highly expressed in distant PCa metastases. We also confirmed that high ALDH1A1 expression in primary tumors predicts disease recurrence indicating its relevance at early stages of prostate cancer progression and suggesting its significance as prognostic biomarker. Consistent with our observations, other studies showed that high ALDH1A1 expression correlates with other prognostic factors such as advanced Gleason score, increased PSA level and pathologic stage, and poor prognosis (Kalantari et al., 2017; Li et al., 2010; Nastały et al., 2019). Contrary to ALDH1A1, high ALDH1A3 expression is predictive for a longer time until progression to castration resistance for patients taking adjuvant hormonal therapy (Wang et al., 2020). Although several ALDH inhibitors have been developed, unfortunately most of them lack clinical viability due to high toxicity, low selectivity and limited efficacy. Improving the anti-cancer efficacy and decreasing normal tissue toxicity of ALDH gene-specific inhibitors, as well as identification of suitable therapy combinations, might open a novel approach for clinical translational studies (Dinavahi et al., 2019).

Our study, for the first time, shows that ALDH1A1 positively regulates TGFB1 expression while ALDH1A3 suppresses it. TGFB1 plays a dual role in PCa development. At the early stages of cancer progression, it acts as a tumor suppressor, whereas in later stages, it plays a role of a tumor promoter by stimulating proliferation, invasion and metastasis (Ahel et al., 2019). Baselga and co-authors observed elevated TGFβ1 levels in patients with advanced disease and skeletal metastasis (Baselga et al., 2008). TGFβ modulates downstream responses via the phosphorylation of the cytoplasmic signaling SMAD proteins (Cao and Kyprianou, 2015). However, in addition to the canonical SMAD signalling, TGFβ acts through the non-canonical SMAD pathway in a context-dependent manner (Zhang, 2009). Remarkably, our data demonstrate that ALDH1A1 and ALDH1A3 knockdown retain the expression of SMAD3, which led us to the conclusion that potential regulation emerges through the non-canonical TGFβ signalling. This might indicate potential mechanisms for the ALDH-dependent regulation of bone marrow metastasis that warrant further investigation.

Recent study by Miftakhova and colleagues demonstrated that ALDH^high^ subpopulation of PC3M cells promotes bone metastatic dissemination by recruiting MMP9 from the host bone marrow (Miftakhova et al., 2016). Our bioinformatics analysis reveals an interconnection of ALDH genes with MMPs on the correlation level. This might suggest additional mechanisms for the ALDH-dependent regulation of bone metastasis that warrant further studies. Recent bioinformatics study of Geng and co-authors revealed MMP11 and MMP26 expression as predictive factors of PCa patient’s recurrence in four different cohorts (Geng et al., 2020). The potential role of MMP genes as biomarkers and prognosticators should be further validated on the cohort of PCa patients. Additional functional validation of our findings is necessary as MMP family members are associated with extracellular matrix degradation and unlock a way for cancer cell spread.

Overall, our findings indicate that ALDH1A1 and ALDH1A3 modulate PCa radiosensitivity and regulate CSCs phenotype and spread of PCa cells to the bone. Our study might have clinical implication for identifying patients at high risk for progression to metastatic disease.

## Acknowledgments

We would like to thank Dr. Power (University of New South Wales, Australia) for sharing the RM1(BM) cell line. We also thank Ellen Geibelt and the Light Microscopy Facility of the Center for Molecular and Cellular Bioengineering in Dresden for assistance with the immunofluorescence analyses. Work in AD lab was partially supported by grants from Deutsche Forschungsgemeinschaft (DFG) (SPP 2084: μBONE, 401326337 and 416001651).

## Abbreviations

ALDH: aldehyde dehydrogenase
AR: androgen receptor
BAAA: bodipy-aminoacetaldehyde
BAA: bodipy-aminoacetate
BM: bone metastatic
CSC: cancer stem cell
DEAB: diethylaminobenzaldehyde
DHT: dihydrotestosterone
DMEM: Dulbecco’s Modified Eagles Medium
DMSO: dimethyl sulfoxide
DoC: duct of Cuvier
dpf: days post fertilization
DTT: dithiothreitol
ECM: extracellular matrix
EGF: epidermal growth factor
EMT: epithelial-mesenchymal transition
Enza: Enzalutamide
FBS: fetal bovine serum
FFPE: fresh frozen paraffin embedded
FGF: fibroblast growth factor
FITC: fluorescein isothiocyanate
FSC: forward scatter
gDNA: genomic DNA
i.c.: intracardiac
IHC: immunohistochemistry
iPSA: initial prostate-specific antigen levels
LN: lymph node
LM: lung metastatic
Luc: luciferase
mCRPC: metastatic castration resistant prostate cancer
MMP: matrix metalloprotease
NSG: NOD scid gamma mouse
P: parental
PBS: phosphate-buffered saline
PCa: prostate cancer
PE: plating efficacy
PFA: paraformaldehyde
PI: propidium iodide
PT: primary tumor
RPMI: Roswell Park Memorial Institute Medium
RR: radioresistant
RT: room temperature
RT: radiation therapy
-RT: sample without added reverse transcriptase enzyme
SF: surviving fraction
shNA: non-silencing small hairpin RNA
shRNA: small hairpin RNA
siRNA: small interfering RNA
siSCR: scrambled small interfering RNA
SPSS: Statistical Package for the Social Sciences
SSC: side scatter
SUMO: Statistical Utility for Microarray and Omics data software
TCGA: The Cancer Genome Atlas
TMA: tissue microarray
TNM: tumor, nodus и metastasis
WF: wide field
Zol: zoledronic acid

## Notes

### Competing Interest Statement

The authors have declared no competing interest.

