## Supplemental Tables 1,2 and Supplemental Figures S1-S8 for "The distinct role of ALDH1A1 and ALDH1A3 in the regulation of prostate cancer metastases"

**Supplementary tables:**
**Table S1.** shRNA constructs used for knockdown of Aldh1a1 and Aldh1a3.
**Table S2.** siRNA oligos, primers and antibodies used in the study.

Table S 1. shRNA constructs used for knockdown of Aldh1a1 and Aldh1a3.

| **Gene** | **Name** | **Vector** | **Hairpin sequence** |
|---|---|---|---|
| Non-silencing | shNS | pLKO.1 | CCTAAGGTTAAGTCGCCCTCGCTC GAGCGAGGGCGACTTAACCTTAGG |
| Aldh1a1 | shAldh1a1 | pLKO.1 | CCCAGTTCTTATCCAAGAATACTC GAGTATTCTTCGGATAAGAACTGGG |
| Aldh1a3 | shAldh1a3 | pLKO.1 | CGAATCCAAGAGTGGAAGAAACTC GAGTTTCTTCCACTCTTGGATTCG |

Table S2. siRNA oligos, primers and antibodies used in the study.

| **siRNA oligos used for knockdown of gene expression** | | |
|---|---|---|
| **Gene** | **siRNA name** | **Sequence** |
| Scrambled siRNA | siSCR | ss GCAGCUAUAUGAAUGUUGUdTdT |
| | | as ACAACAUUCAUAUAGCUGCdTdT |
| ALDH1A1 | siALDH1A1 #1 | ss GAUCCAGGGCCGUACAAUAdTdT |
| | | as UAUUGUACGGCCCUGGAUCdTdT |
| ALDH1A1 | siALDH1A1 #2 | ss AGGUAGAAGAAGGAGAUAAdTdT |
| | | as ACAACAUUCAUAUAGCUGCdTdT |
| ALDH1A3 | siALDH1A3 #1 | ss AGGAAAUGGCAGAGAACUAdTdT |
| | | as UAGUUCUCUGCCAUUUCCUdTdT |
| ALDH1A3 | siALDH1A3 #2 | ss UCGUGGAGGAGCAGGUCUAdTdT |
| | | as UAGACCUGCUCCUCCACGAdTdT |
| Scrambled | siSCR | ss GCAGCUAUAUGAAUGUUGUdTdT |
| siRNA pool | | as ACAACAUUCAUAUAGCUGCdTdT |
| | | ss UGCGCUAGGCCUCGGUUGCdTdT |
| | | as GCAACCGAGGCCUAGCGCAdTdT |
| AR | siAR | ss CCAAAGGGCUAGAAGGCGAdTdT |
| | | as UCGCCUUCUAGCCCUUUGGdTdT |
| | | ss AUUGAUAAAUUCCGAAGGAdTdT |
| | | as UCCUUCGGAAUUUAUCAAUdTdT |
| CTNNB1 | siCTNNB1 | ss GGGUACGAGCUGCUAUGUUdTdT |
| | | as AACAUAGCAGCUCGUACCCdTdC |
| | | ss GGUGGUGGUUAAUAAGGCUdTdT |
| | | as AGCCUUAUUAACCACCACCdTdG |

| **Primers used for RT-qPCR** | |
|---|---|
| **Gene** | **Sequence (5’→3’)** |
| ACTB | F 5'- ATGGAGTCCTGTGGCATCCA -3' |
| | R 5'- AGTACTTGCGCTCAGGAGGA -3' |
| RPLP0 | F 5'- CTCAACATCTCCCCCTTCTCCTT -3' |
| | R 5'- TGATGCAACAGTTGGGTAGCC -3' |
| ALDH1A1 | F 5'- GAATGGCATGATTCAGTGAGTGG-3' |
| | R 5'- CAGCCAACTTGTATAATAGTCG-3' |
| ALDH1A3 | F 5'- TCTCGACAAAGCCCTGAAGT-3' |

| | |
|---|---|
| | R 5'- TATTCGGCCAAAGCGTATTC -3' |
| AR | F 5'- CATCTTGTCGTCTTCGGAAATGTTA -3' |
| | R 5'- GAAGCCTCTCCTTCCTCCTGTAGTT -3' |
| CTNNB1 | F 5'- ATTTGATGGAGTTGGACATGGC-3' |
| | R 5'- CCAGCTACTTGTTCTTGAGTGAAGG -3' |
| SMAD3 | F 5'- GCGTGCGGGTCTACTACATC -3' |
| | R 5'- GCACATTCGGGTCAACTGGTA -3' |
| TGFB1 | F 5'- CAGCAGGGATAACACACTGC -3' |
| | R 5'- CACGCAGCAGTTCTTCTCC -3' |
| Mouse/rat Gapdh | F 5'- TTCAACGGCACAGTCAAGG -3' |
| | R 5'- ACATACTCAGCACCAGCATCAC -3' |
| Mouse Aldh1a1 | F 5'- ATACTTGTCGGATTTAGGAGGCT-3' |
| | R 5'- GGGCCTATCTTCCAAATGAACA -3' |
| Mouse Aldh1a3 | F 5'- GGGTCACACTGGAGCTAGGA -3' |
| | R 5'- CTGGCCTCTTCTTGGCGAA -3' |

**Antibodies used for Immunofluorescence**

| **Antibody** | **Dilution and host** | **Vendor and catalogue number** |
|---|---|---|
| Endomucin | 1:200, Goat | Thermo Fisher Scientific, #PA5-47648 |
| GFP Tag | 1:200, Rabbit | Thermo Fisher Scientific, #A6455 |
| Anti-Goat IgG (Alexa Fluor 555) | 1:350, Donkey | |
| Anti-Rabbit IgG (Alexa Fluor 488) | 1:350, Donkey | |

**Supplementary Figures:**

**Figure S1.** ALDH1A1 and ALDH1A3 as regulators of radiosensitivity.

**Figure S2.** Correlation analysis of ALDH1A1 or ALDH1A3 expression in the TCGA PRAD dataset with RT2 signatures.

**Figure S3.** qPCR validation of 84 extracellular matrix and adhesion genes analysed using DNA RT² Profiler PCR Array.

**Figure S4.** Disease-free survival of prostate cancer patients with high or low ALDH1A1 and ALDH1A3 expression.

**Figure S5.** Survival and extravasation potential of color-coded PC3 cells.

**Figure S6.** Comparison of the iPSA serum level and nuclear AR between tumors with or without ALDH1A1 or ALDH1A3 expression.

**Figure S7.** Analysis of the osteogenesis-related genes.

**Figure S8.** Interconnection of ALDH genes with MMPs.

### Supplementary Figure S1

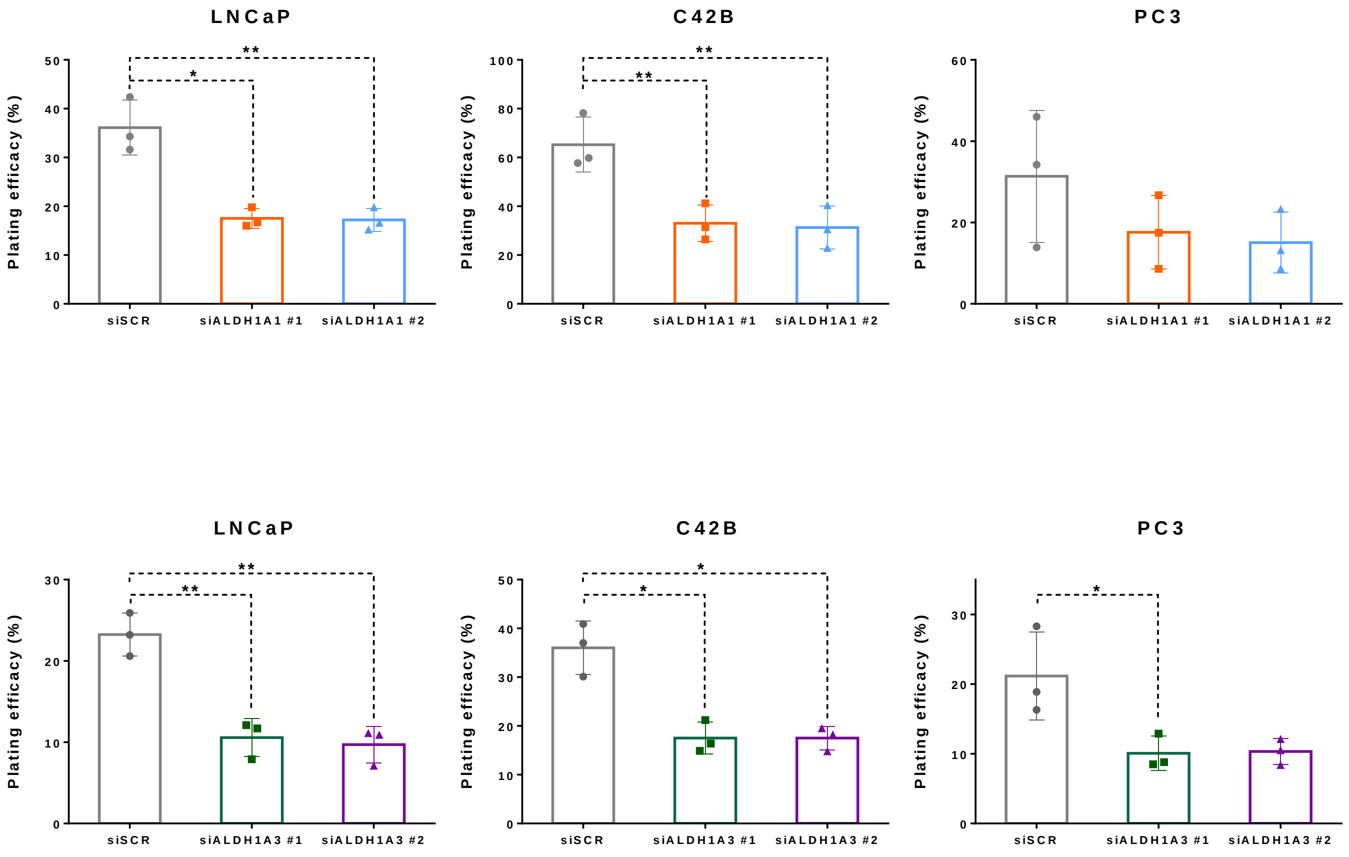

### Supplementary Figure S2

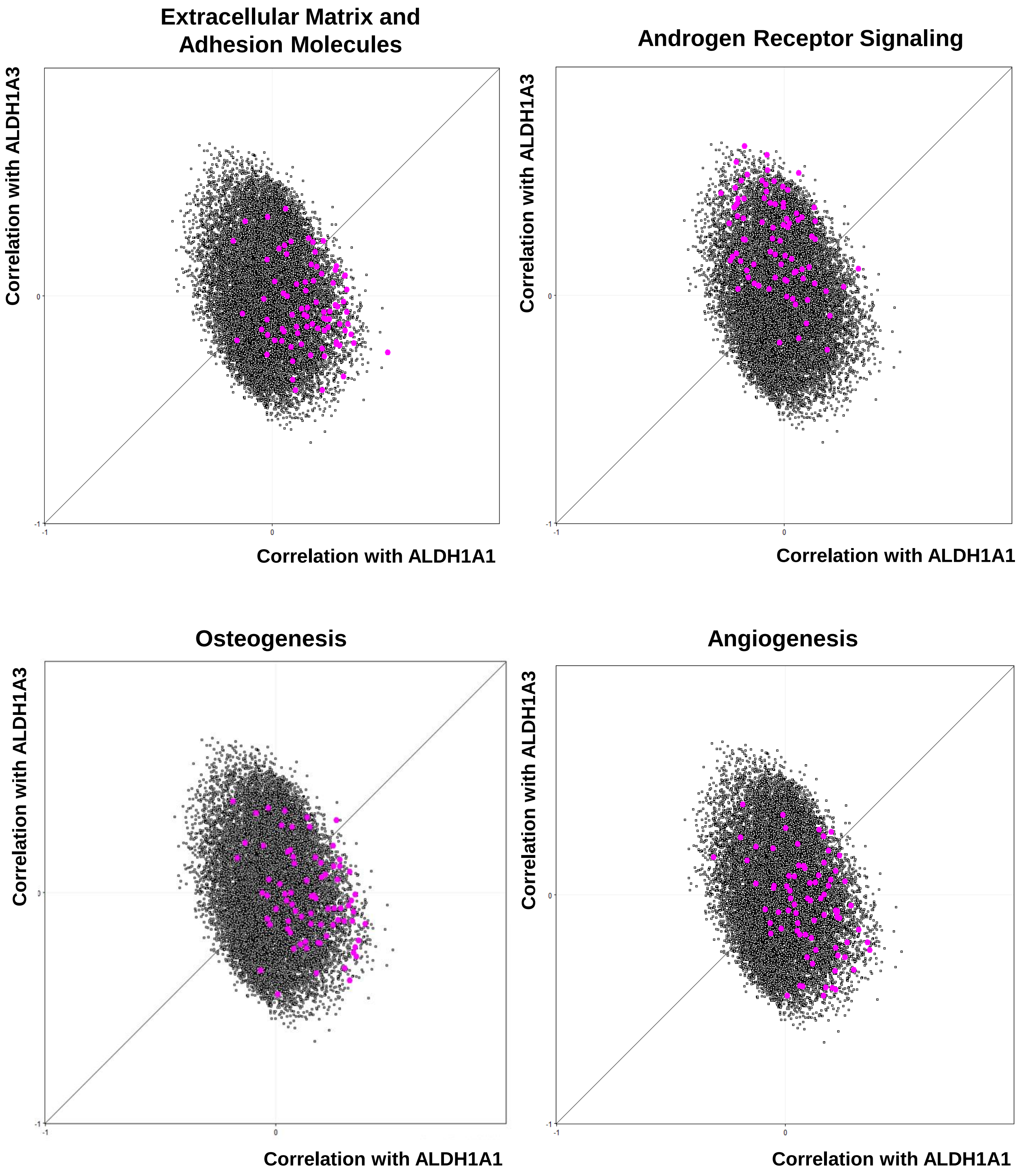

### Supplementary Figure S3

| | LNCaP | | | | PC3 | |
|---|---|---|---|---|---|---|
| | siALDH1A1 | siALDH1A3 | | | siALDH1A1 | siALDH1A3 |
| ITGA8 | 3,4457424 | 13,176562 | | ADAMTS1 | 2,5456312 | 1,2379519 |
| ITGA1 | 3,3190363 | 2,5670124 | | ADAMTS13 | 2,4899838 | 1,7253882 |
| SELE | 2,8402345 | 1,9934559 | | COL7A1 | 2,4816593 | 3,2576491 |
| LAMB1 | 1,8026658 | 1,7948206 | | COL16A1 | 2,4191095 | 3,4794358 |
| MMP8 | 1,7800653 | 1,8855528 | | VTN | 1,9934757 | 0,9320517 |
| COL14A1 | 1,6748544 | 3,5283427 | | ITGAM | 1,9313438 | 1,3742763 |
| VTN | 1,6677947 | 1,6008062 | | COL14A1 | 1,91886 | 1,7008488 |
| ITGB3 | 1,597696 | 2,1582947 | | VCAN | 1,9041017 | 0,9428615 |
| MMP3 | 1,5931042 | 0,7725707 | | COL12A1 | 1,7694796 | 1,2398371 |
| MMP10 | 1,5807678 | 0,3869222 | | LAMB1 | 1,7525492 | 1,0384546 |
| MMP9 | 1,4836756 | 0,9496065 | | ITGA7 | 1,6752365 | 1,5290232 |
| ITGA2 | 1,3291003 | 0,6567818 | | LAMA3 | 1,6398663 | 1,1025826 |
| PECAM1 | 1,2995651 | 1,7689802 | | LAMA1 | 1,553781 | 0,3770434 |
| ICAM1 | 1,2680028 | 1,0782503 | | MMP7 | 1,5237628 | 0,733172 |
| COL4A2 | 1,2524879 | 1,2067284 | | TIMP2 | 1,5073309 | 0,5509773 |
| THBS3 | 1,1880757 | 0,8636981 | | THBS3 | 1,4554457 | 1,4294116 |
| SPG7 | 1,1627652 | 1,9019644 | | ICAM1 | 1,4349864 | 1,1119441 |
| COL7A1 | 1,1585684 | 1,2383869 | | MMP12 | 1,4180188 | 1,7192085 |
| ITGB1 | 1,1448859 | 1,14006 | | ITGAV | 1,4154902 | 1,4296441 |
| ADAMTS13 | 1,1315893 | 1,1552432 | | VCAM1 | 1,4075145 | 3,412206 |
| MMP13 | 1,122938 | 0,6891376 | | LAMC1 | 1,3905187 | 1,1085608 |
| TGFBI | 1,0978999 | 0,8807619 | | MMP15 | 1,3893775 | 0,7165384 |
| COL1A1 | 1,0977011 | 0,9863823 | | THBS2 | 1,3845623 | 1,662503 |
| ITGA4 | 1,0878769 | 0,2124813 | | SPG7 | 1,3782269 | 1,009498 |
| LAMC1 | 1,0401072 | 1,0576065 | | SELE | 1,33 | 0,7924925 |
| TIMP2 | 1,034769 | 0,7548247 | | TIMP1 | 1,3329303 | 0,6703062 |
| LAMA3 | 1,0317555 | 1,8088347 | | ITGA2 | 1,3112203 | 0,8404221 |
| ADAMTS8 | 1,0021507 | 1,4579233 | | HAS1 | 1,2904627 | 0,9911907 |
| ITGAV | 0,9775376 | 1,0227054 | | MMP16 | 1,2864494 | 1,5395976 |
| NCAM1 | 0,9751716 | 1,1230701 | | CLEC38 | 1,2562167 | 1,2960533 |
| COL6A2 | 0,9688324 | 1,1562884 | | ITGA5 | 1,2492804 | 1,3850067 |
| ITGA7 | 0,9675012 | 0,851476 | | ECM1 | 1,1973863 | 2,011028 |
| COL16A1 | 0,9626171 | 1,1075554 | | SGCE | 1,1905576 | 0,8772027 |
| MMP12 | 0,9606135 | 0,8372479 | | CTNNB1 | 1,161265 | 1,0290504 |
| CDH1 | 0,9400942 | 1,0851935 | | ITGB4 | 1,1529279 | 1,3502799 |
| MMP16 | 0,9373515 | 2,9326135 | | ADAMTS8 | 1,1280559 | 1,2660885 |
| ADAMTS1 | 0,9283781 | 1,147903 | | ITGA4 | 1,0983899 | 1,6256578 |
| MMP15 | 0,9188098 | 1,2483633 | | COL1A1 | 1,090569 | 2,1716679 |
| CTNNB1 | 0,9066081 | 0,9445517 | | CCN2 | 1,0466456 | 0,6457103 |
| COL12A1 | 0,8844819 | 1,4549272 | | COL4A2 | 1,0433044 | 0,8014654 |
| ITGAL | 0,8809342 | 0,8944822 | | COL5A1 | 1,0384563 | 0,7542075 |
| THBS1 | 0,8725112 | 2,3957273 | | ITGB1 | 1,0339793 | 0,9750346 |
| ECM1 | 0,869554 | 1,8347866 | | LAMA2 | 1,0326569 | 1,283846 |
| ITGA5 | 0,8644463 | 1,0651375 | | CNTN1 | 1,0259969 | 1,6086714 |
| MMP11 | 0,8582322 | 0,7557293 | | COL6A1 | 1,0210848 | 1,4432137 |
| HAS1 | 0,8576526 | 0,7661647 | | SELL | 1,01 | 1,4612911 |
| LAMB3 | 0,8434573 | 2,437513 | | MMP14 | 1,0027862 | 0,6310447 |
| CTNND1 | 0,8338764 | 1,1585378 | | CTNND1 | 0,9969843 | 0,9894442 |
| ITGA6 | 0,8330478 | 0,9970363 | | ITGA3 | 0,9852726 | 1,5170697 |
| SGCE | 0,8213126 | 0,9780573 | | CCNNA1 | 0,9794878 | 1,7519162 |
| CCNNA1 | 0,8209316 | 1,0206037 | | MMP9 | 0,968892 | 0,9657469 |
| ITGB5 | 0,8047896 | 0,7803178 | | MMP11 | 0,9649789 | 1,5369416 |
| LAMA2 | 0,8009228 | 0,9926408 | | TGFBI | 0,9321974 | 0,8030554 |
| COL6A1 | 0,7879602 | 1,0667271 | | COL6A2 | 0,9293094 | 0,9736498 |
| MMP7 | 0,7840893 | 0,7566001 | | ITGB5 | 0,9088361 | 0,6444732 |
| MMP1 | 0,7831983 | 0,6783706 | | SPP1 | 0,8934002 | 0,460759 |
| ITGA3 | 0,7761631 | 1,7270719 | | CDH1 | 0,8517873 | 5,0105759 |
| FN1 | 0,7647732 | 1,0024687 | | MMP3 | 0,8382109 | 0,4833455 |
| COL5A1 | 0,7641728 | 0,7017601 | | ITGA6 | 0,8282865 | 1,1035918 |
| ITGB4 | 0,6895869 | 1,0211152 | | ITGB3 | 0,8094661 | 0,3686243 |
| LAMA1 | 0,6640285 | 0,4835741 | | MMP10 | 0,7854759 | 0,4220869 |
| ITGB2 | 0,6415707 | 1,0311173 | | THBS1 | 0,7723966 | 1,8800374 |
| SELL | 0,6392187 | 1,3916102 | | CD44 | 0,7711538 | 1,2881271 |
| TIMP3 | 0,6117086 | 1,272633 | | FN1 | 0,7458408 | 0,686763 |
| CLEC38 | 0,5540223 | 0,7528494 | | TNC | 0,720417 | 0,6379045 |
| CCN2 | 0,4157619 | 0,6769479 | | ITGA1 | 0,7046165 | 1,0064237 |
| TNC | 0,4044734 | 0,7966553 | | ITGB2 | 0,6361033 | 1,3870517 |
| SPP1 | 0,336207 | 0,3857541 | | CTNND2 | 0,6086106 | 0,8425904 |
| SPARC | 0,2264424 | 0,606537 | | ANOS1 | 0,4683601 | 0,6615725 |
| | | | | MMP13 | 0,4479699 | 0,518483 |
| | | | | COL8A1 | 0,3564923 | 1,4662142 |
| | | | | MMP1 | 0,3429869 | 0,3899497 |
| | | | | PECAM1 | 0,33 | 0,6189817 |
| | | | | SPARC | 0,2649995 | 0,5830115 |

### Supplementary Figure S4

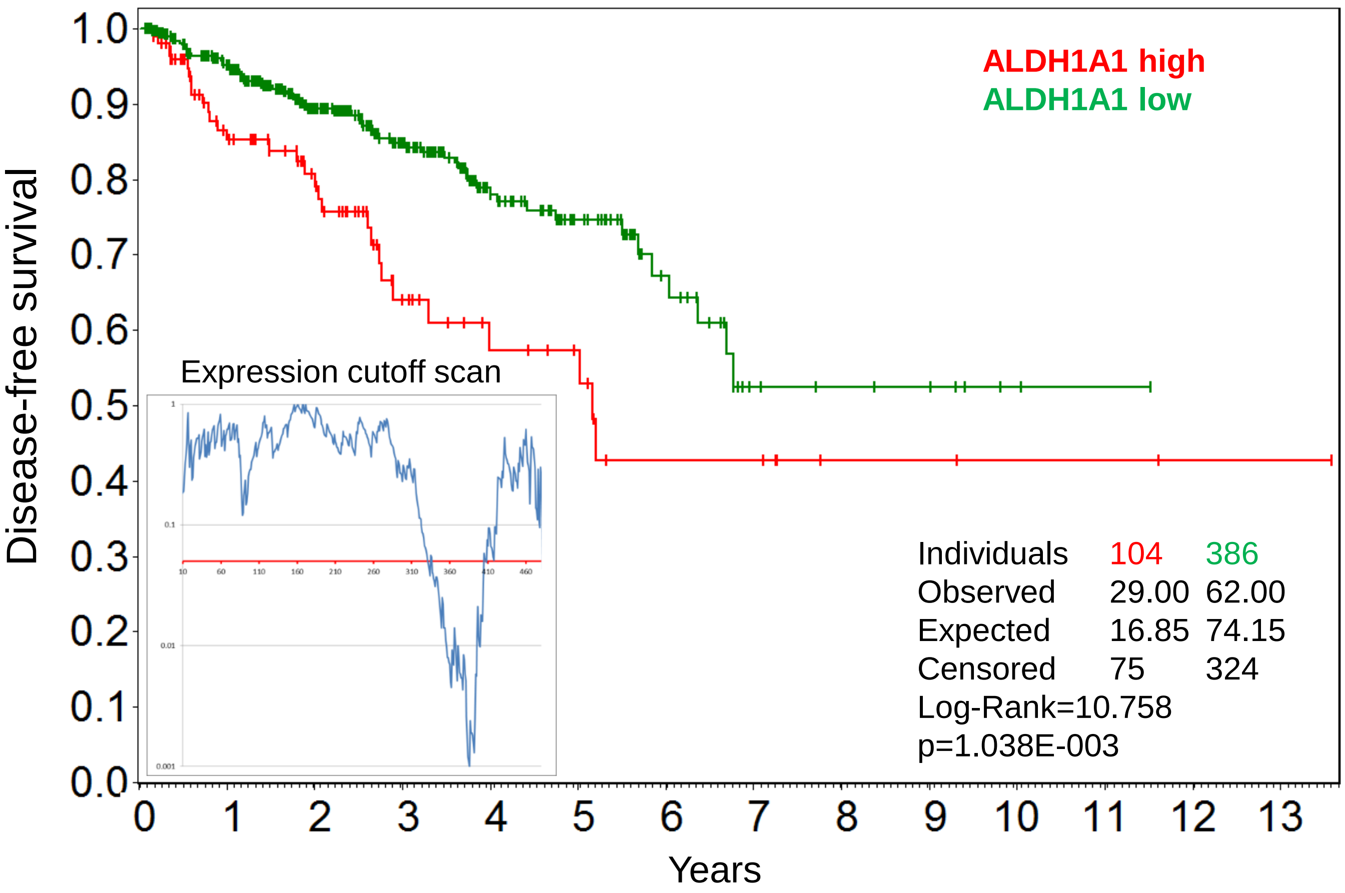


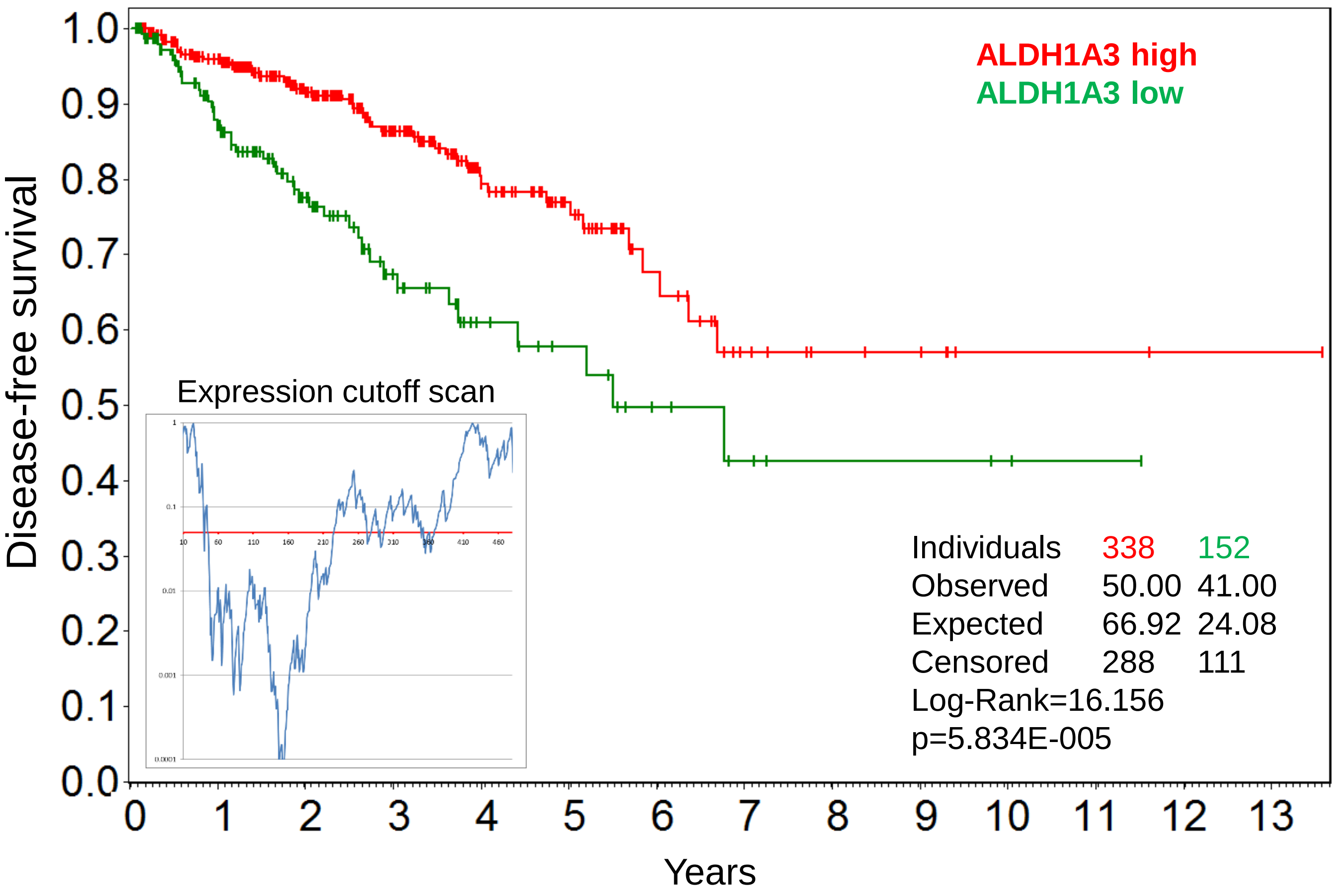

### Supplementary Figure S5

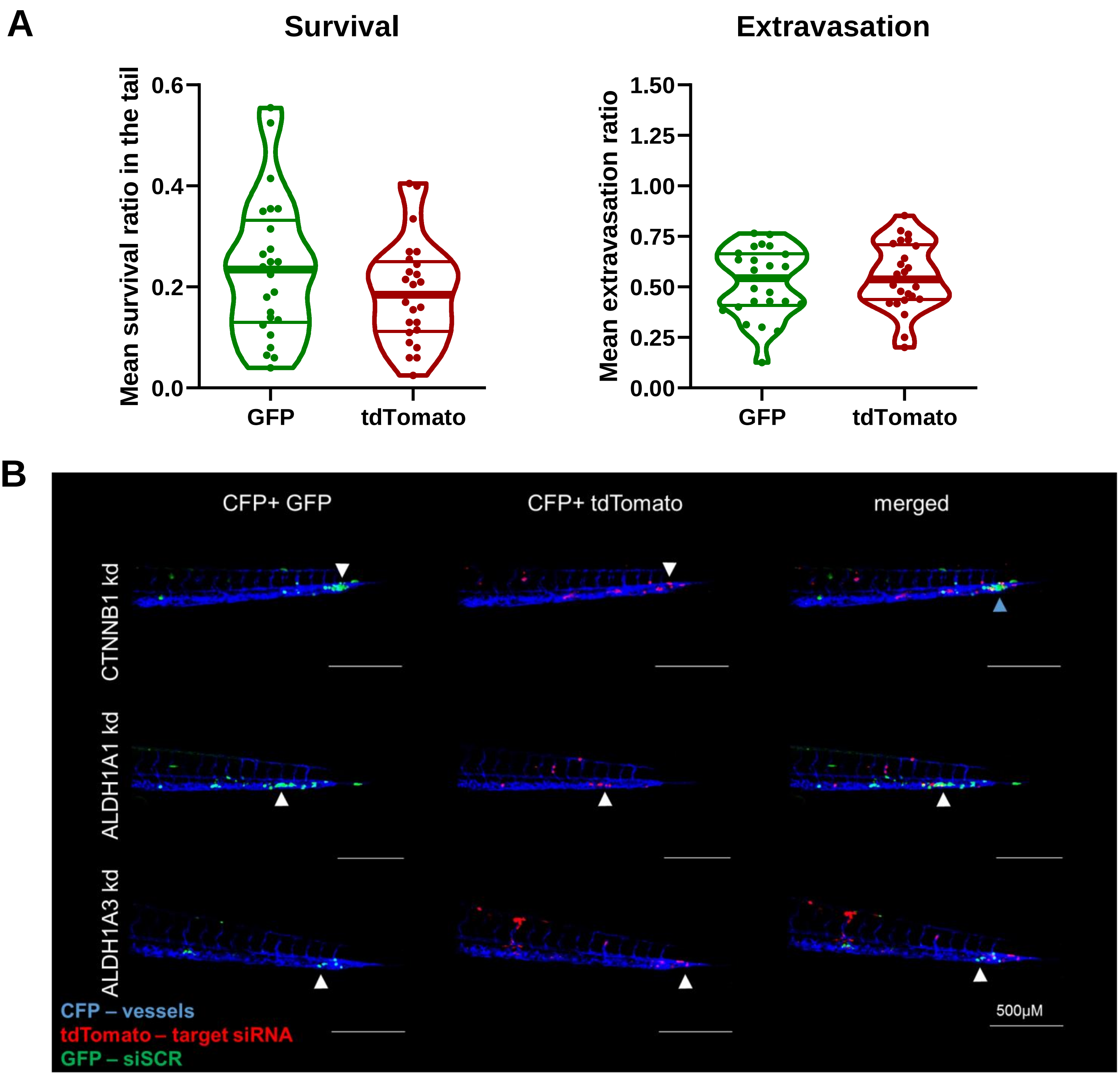

### Supplementary Figure S6

**A**

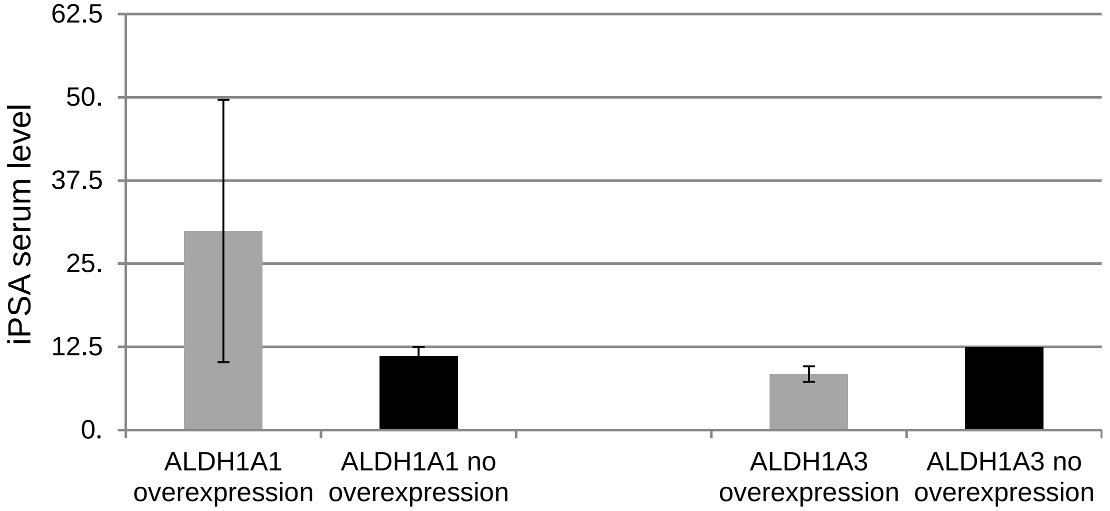


**B**

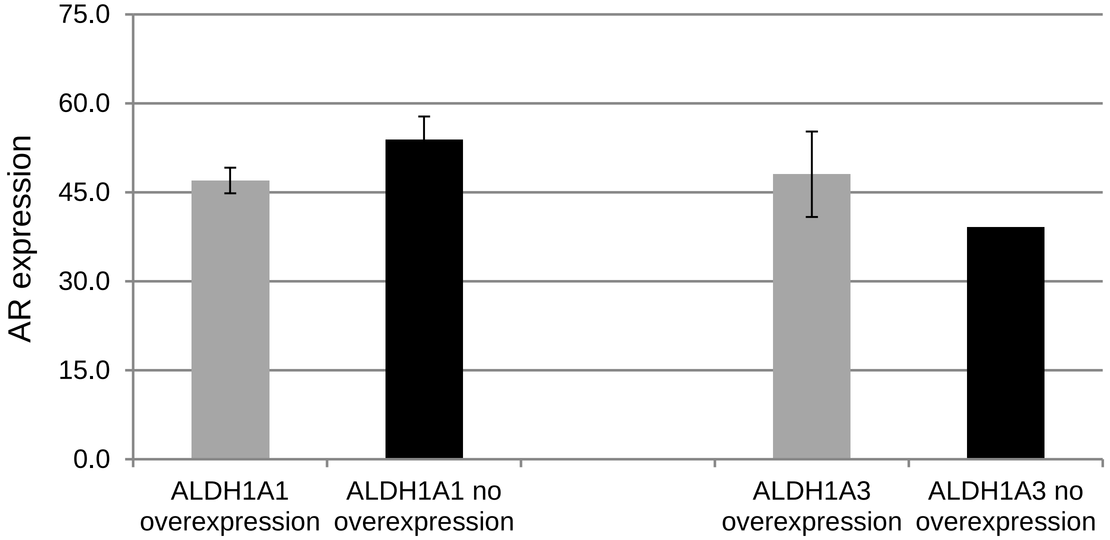

### Supplementary Figure S7

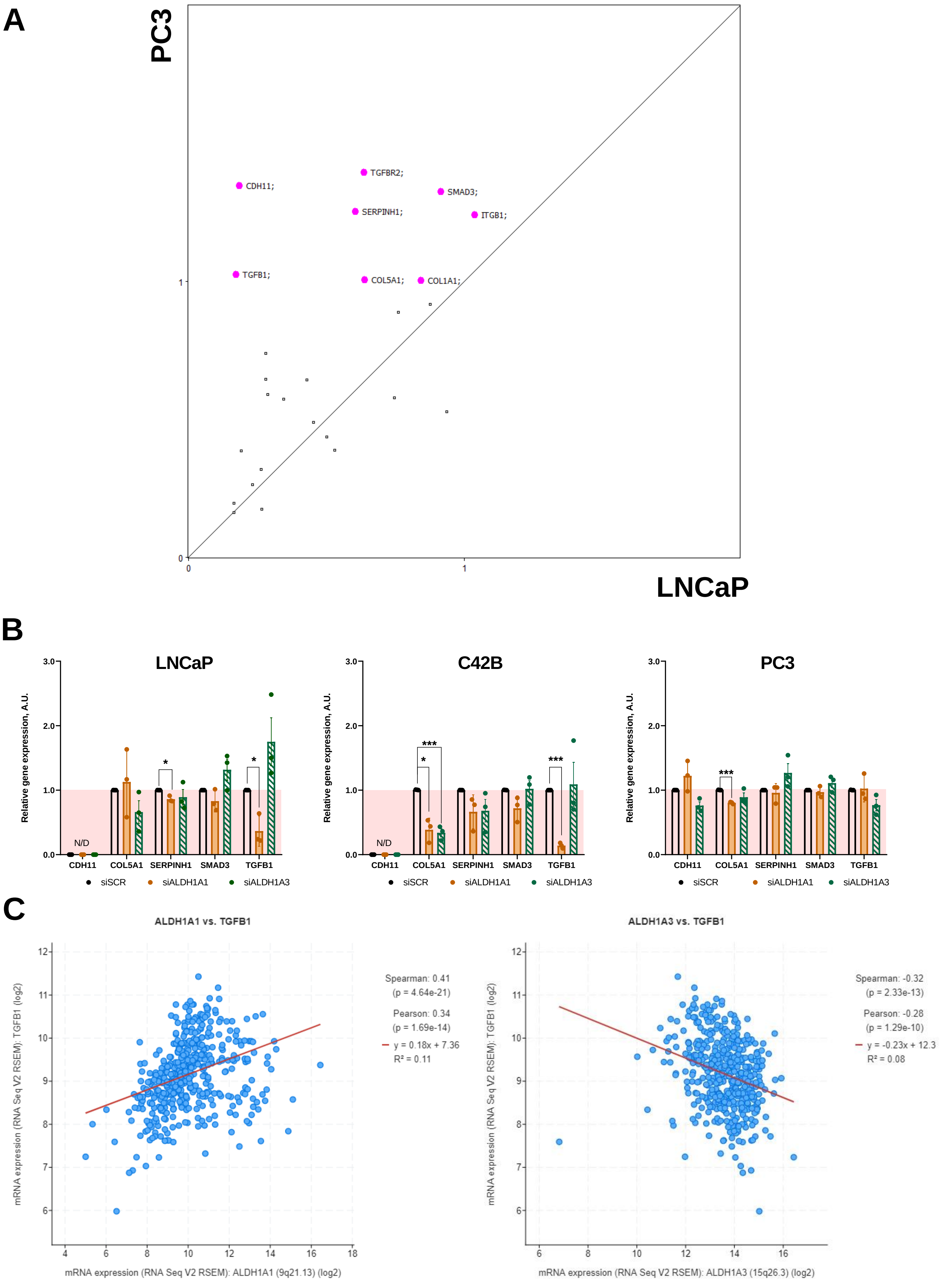

### Supplementary Figure S8

**A**

| LNCaP | ALDH1A1kd | ALDH1A3kd |
|---|---|---|
| MMP1 | 0,783198275 | 0,678370572 |
| MMP7 | 0,784089265 | 0,756600085 |
| MMP11 | 0,858232194 | 0,755729342 |
| MMP15 | 0,918809791 | 1,248363293 |
| MMP16 | 0,937351456 | 2,932613542 |
| MMP12 | 0,960613538 | 0,837247884 |
| MMP13 | 1,122938011 | 0,689137573 |
| MMP9 | 1,483675634 | 0,949606482 |
| MMP10 | 1,580767792 | 0,386922226 |
| MMP3 | 1,593104239 | 0,772570745 |
| MMP8 | 1,780065293 | 1,885552818 |

| PC3 | ALDH1A1kd | ALDH1A3kd |
|---|---|---|
| MMP1 | 0,3429869 | 0,38994966 |
| MMP13 | 0,44796986 | 0,51848302 |
| MMP10 | 0,78547594 | 0,42208688 |
| MMP3 | 0,83821089 | 0,48334545 |
| MMP11 | 0,9649789 | 1,5369416 |
| MMP9 | 0,96889195 | 0,9657469 |
| MMP14 | 1,00278618 | 0,63104465 |
| MMP16 | 1,28644942 | 1,53959762 |
| MMP15 | 1,38937753 | 0,7165384 |
| MMP12 | 1,41801883 | 1,71920845 |
| MMP7 | 1,52376282 | 0,73317204 |

**B**

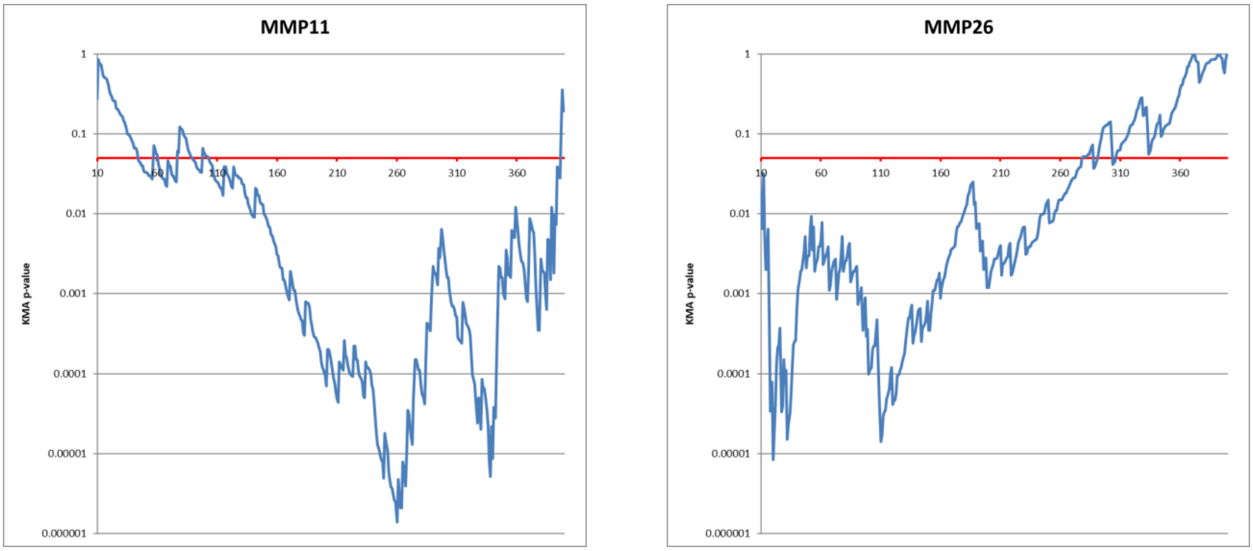


**C**

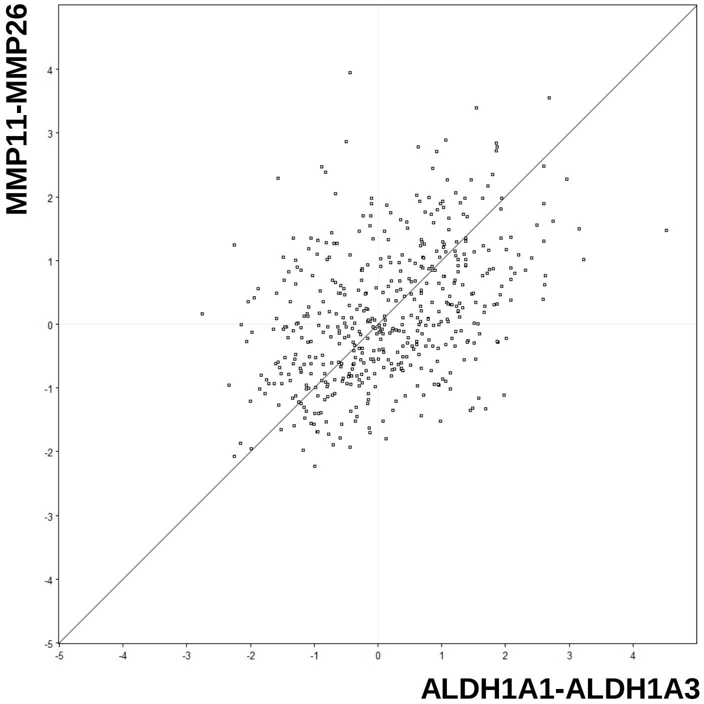


Pearson correlation coefficient:
ALDH1A1-ALDH1A3 / MMP11-MMP26;
n=490;r=0.42670; p=4.23E-023
